# Microbiome-mediated lauric acid turnover contributes to colonization resistance against *Ralstonia* in the tomato rhizosphere

**DOI:** 10.64898/2026.05.12.723958

**Authors:** Shengyue Tang, Hao Li, Pugang Yu, Qingliu Wu, Ting Xiao, Yuxiao Huang, Fusuo Zhang, Bin Ni

## Abstract

Host-associated lipids shape plant–microbiome interactions, but their microbiome-mediated turnover and role in pathogen exclusion are not fully understood. In tomato, lauric acid (C12:0) occurs across root-associated compartments, and its abundance at the root–soil interface is linked to soil-dependent differences in rhizosphere community composition and *Ralstonia solanacearum* colonization. Lauric acid promotes *Ralstonia* motility and virulence in *vitro* while inhibiting some beneficial lauric-acid-sensitive taxa, especially Gram-positive antagonists. In disease-suppressive soils, rhizosphere communities enriched in lauric-acid-degrading taxa are associated with lower local lauric acid levels and reduced pathogen colonization, suggesting community-dependent buffering of the rhizosphere lauric acid pool alongside other protective microbiome functions. Soil perturbation, microbiome transplantation, gnotobiotic SynCom reconstruction, and isotope tracing provide convergent evidence that microbiome-mediated lauric acid turnover contributes to colonization resistance and helps explain how rhizosphere lipid chemistry influences pathogen invasion outcomes in tomato, revealing an ecologically grounded protective mechanism for pathogen management.

## Introduction

Plants exist within a complex ecological niche in which their growth, development, and immunity are profoundly influenced by microbial communities inhabiting the rhizosphere—the narrow soil zone shaped by root exudates. This rhizosphere microbiota encompasses a taxonomically and functionally diverse assemblage of bacteria, fungi, archaea, viruses, and protists that collectively extend the plant’s physiological capacity by modulating nutrient uptake, pathogen resistance, and abiotic stress responses^1–5^. Far from being passive residents, these microbes form dynamic, structured consortia characterized by metabolic interdependence, niche partition and ecological specialization^6–12^. Their composition and functionality are sculpted by a network of interacting factors, including host genotype, developmental stage, soil physicochemical properties, and agricultural practices^13–17^. Unraveling the molecular logic by which plants recruit, assemble, and sustain beneficial microbiomes remains a central challenge in plant biology with significant implications for sustainable agriculture.

At the heart of plant–microbiome communication lies the chemical interface of root exudates. These secretions—ranging from simple sugars and amino acids to complex lipids and secondary metabolites—serve dual roles as nutrient sources and signaling molecules that mediate both host–microbe and microbe–microbe interactions^6,18,19^. Their chemical diversity enables selective microbial recruitment or inhibition. Hydrophilic compounds such as organic acids and amino acids diffuse readily into the surrounding soil, acting as chemoattractants and carbon sources for mutualistic microbes, including nitrogen-fixing rhizobia and phosphate-solubilizing bacteria^20–22^. In contrast, hydrophobic exudates—such as fatty acids, oxylipins, and cutin monomers—remain localized at the root–soil interface, influencing microbial adhesion, niche partitioning, and biofilm formation^23^. The physicochemical properties of these compounds dictate their spatial reach and bioavailability, revealing a nuanced chemical ecology within the rhizosphere. Many plant species secrete fatty acids as components of their root exudates, contributing to critical processes such as plant-microbe interactions. Although comprehensive profiling across plant families is limited, notable examples of fatty acid exudation have been observed in *Arabidopsis thaliana*, tomato (*Solanum lycopersicum*), and maize (*Zea mays*)^24–26^. Despite advances in understanding sugar- and organic acid-mediated microbial recruitment, the role of root-derived lipids—particularly fatty acids and oxylipins—as regulators of microbial community structure and pathogen resistance remains largely unexplored.

Importantly, the exudate profile is dynamic and responsive to both developmental and environmental cues. Specific microbial colonization patterns are partially governed by host genotype through tailored exudate profiles^6,27^. Under nutrient limitation or biotic stress, plants undergo “exudation reprogramming,” altering the composition and quantity of exudates to recruit protective microbes or suppress deleterious taxa^28–31^. These shifts can trigger positive feedback loops in which recruited microbes metabolize exudates into bioactive compounds that further influence plant immunity, hormone signaling, and root morphology—demonstrating a complex, bidirectional communication axis^32–35^. Root exudates also mediate microbial interactions, fostering cooperation, competition, and signaling. Hydrophilic compounds broadly support microbial proliferation, while hydrophobic exudates help spatially structure microbial communities. Certain exudates further modulate microbial behavior by mimicking or disrupting quorum-sensing signals^23,36–38^.

Lipids are emerging as potent, yet underexplored, regulators of rhizosphere microbiome composition and function. Although traditionally recognized as dietary components that influence the gut microbiota—affecting processes such as memory, obesity, infection, and immune responses^39–47^—lipids also play critical roles in plant-associated microbiomes. Lipid-derived compounds, including fatty acids, oxylipins, and jasmonates, function not only as carbon sources but also as signaling molecules that influence microbial recruitment, community structure, and pathogen interactions in the rhizosphere^25,48–51^. These molecules can selectively promote the colonization of beneficial taxa while suppressing opportunistic pathogens, effectively acting as molecular gatekeepers of the root microbiome^49,52,53^. Fatty acids, defined by a hydrophilic carboxyl group and a hydrophobic aliphatic tail, are classified by chain length into short-, medium-, and long-chain species. These molecules can act both as antimicrobial agents that shape microbial competition and as signaling mediators that modulate microbial behavior, positioning them as potentially important regulators of rhizosphere microbial ecology^48,54^. Concepts from mammalian lipid–microbiome interactions provide a useful heuristic for thinking about host-associated lipids in the rhizosphere. However, whether plant-associated fatty acids and their microbial turnover similarly structure rhizosphere communities and influence pathogen exclusion remains largely unresolved^55^. In mammals, the immunomodulatory and microbiota-regulatory roles of host-derived lipids are well established; we refer to these systems only as conceptual analogies rather than as evidence for a conserved mechanism in plants^56,57^. For example, oleic acid promotes the proliferation of *Lactobacillus crispatus*, supporting vaginal microbiome stability, while long-chain fatty acids regulate intestinal inflammation and immune tone^47,58–60^. Lauric acid, in particular, exhibits potent and broad-spectrum antimicrobial activity against enveloped viruses and various bacteria, and has been utilized therapeutically in dermatological treatments^61^.

In plants, the rhizosphere microbiome not only supports growth and nutrient acquisition but also constitutes a critical defense layer against pathogens. A prime example is *Ralstonia solanacearum*, a devastating soilborne pathogen that causes bacterial wilt in many crops and exemplifies the complex interplay between pathogen virulence, root exudate chemistry, and protective microbial consortia. Successful infection by *Ralstonia* requires navigating the rhizosphere, breaching root barriers, and colonizing the xylem—a process reliant on motility, chemotaxis toward exudates, and immune evasion through biofilm formation and type III secretion systems^62–65^. However, successful infection of plant hosts by this pathogen is often constrained by the resident rhizosphere microbiota. Beneficial microbes such as *Pseudomonas*, *Bacillus*, and *Streptomyces* inhibit pathogen establishment through competition for niches and nutrients, production of antimicrobial compounds, and activation of plant immune responses^66,67^. These beneficial communities actively remodel the rhizosphere environment to limit pathogen access to entry sites.

Soil type, plant genotype, and agricultural practices jointly shape the composition and activity of microbial communities, often leading to disease-suppressive soils enriched with taxa such as Burkholderiaceae, Sphingomonadaceae, and Actinobacteria—microbial groups recognized for their roles in antimicrobial production, nutrient competition, and quorum quenching^68–71^. Notably, the protective capacity of the rhizosphere microbiome is not solely a function of its taxonomic composition but also of its ecological functionality, including network resilience, functional redundancy, and colonization resistance—the ability of a community to resist pathogen invasion by maintaining structural and metabolic integrity^72^. Recent advances in microbial community profiling, synthetic microbial consortia, theoretical microbial ecology, and plant-microbiome interactions are beginning to reveal the underlying principles that govern these emergent properties.

Despite extensive work on sugar- and organic-acid-mediated host–microbe interactions, the role of plant-associated lipids and their microbial turnover in pathogen exclusion remains poorly understood. Here, we investigate how lauric acid is distributed across tomato root-associated compartments, how it is transformed by rhizosphere communities, and how these processes influence susceptibility to *Ralstonia*. We identify rhizosphere communities enriched in lauric-acid-degrading taxa, quantify spatial variation in lauric acid across soil compartments, and test how microbiome perturbation and reconstruction alter pathogen colonization. Our results support a model in which microbiome-mediated lauric acid turnover contributes to colonization resistance in a soil- and compartment-dependent manner, while acting alongside other microbiome functions that also shape pathogen invasion outcomes. We therefore treat rhizosphere LA concentration as an emergent property of plant-associated input and microbial turnover, rather than as a direct proxy for secretion alone; the relative contributions of host release, physicochemical retention, and microbial turnover remain to be resolved.

## Results

### Microbiome-mediated root colonization resistance against *Ralstonia* in tomato rhizosphere

Tomato plants of the same cultivar were grown in soils collected from 8 representative regions in China (Fig. 1A,F; Fig. S1A). Plant performance varied markedly among soils: tomatoes grown in SD and HLJ soils exhibited robust growth, while those in HB and GX soils showed severe stunting, with reductions of ∼63.0% in dry weight (Fig. S1C), ∼61.8% in fresh weight (Fig. S1D), ∼40.4% in height (Fig. S1E), and ∼35.7% in stem diameter (Fig. S1F). To investigate the microbial and environmental determinants of plant health and susceptibility to bacterial wilt, we profiled the rhizosphere and bulk soil microbiomes. Both bacterial and fungal communities differed significantly across regions (Fig. 1B,C), with bacterial communities showing stronger regional separation by NMDS (Fig. 1D) than fungal communities (Fig. 1E). Compared to fungi, bacterial richness was significantly lower in rhizosphere soils (Fig. S2A,B), suggesting selective recruitment by tomato roots. Specifically, the rhizosphere was enriched in *Pseudomonas*, *Flavobacterium*, and *Arthrobacter*, and depleted in *Sphingomonas*, *Delftia*, and *Bacillus* (Fig. 1B), highlighting functional microbial selection.

**Fig. 1.**
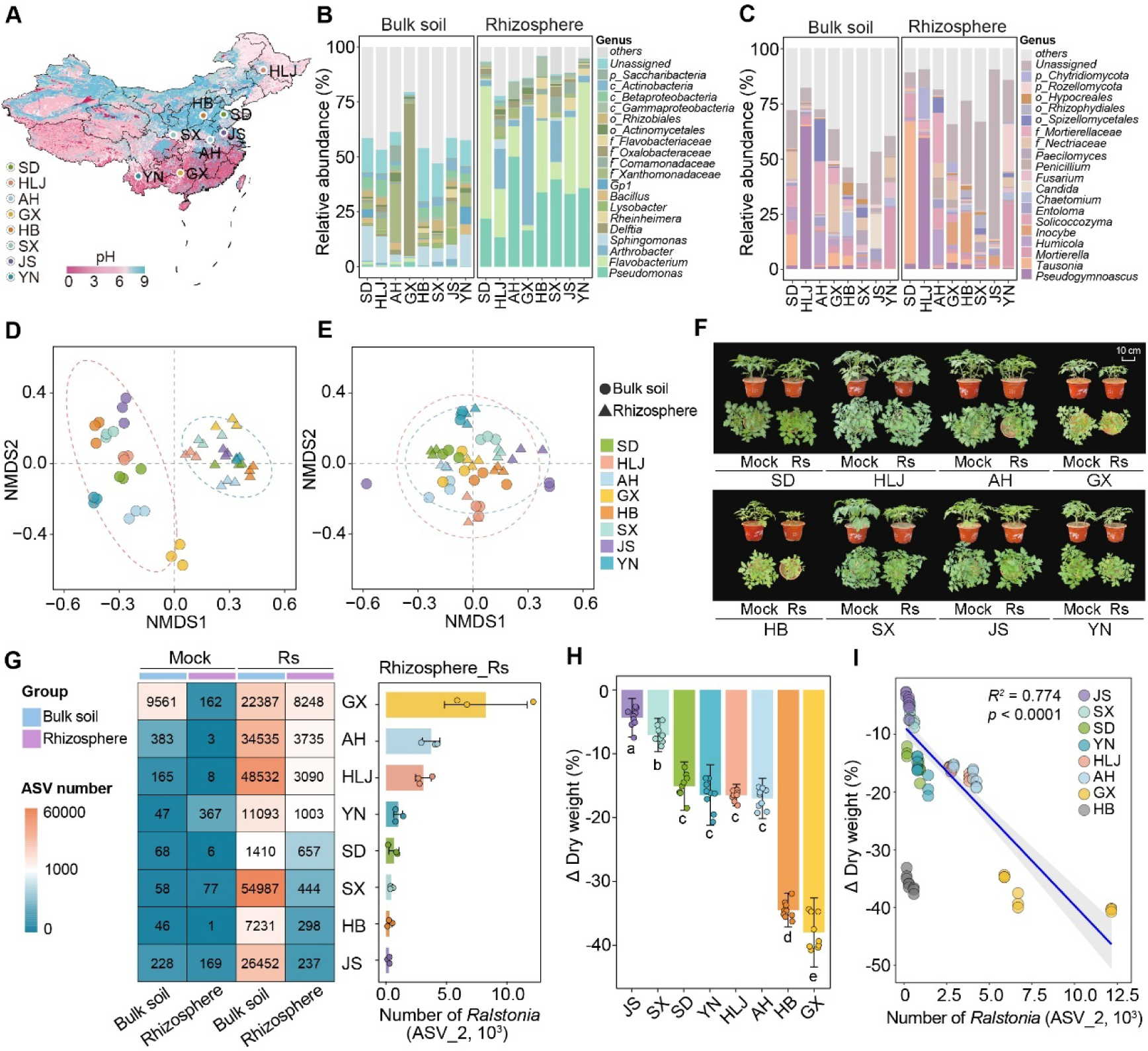
Microbiome-mediated phenotypic plasticity in colonization resistance to pathogenic *Ralstonia* in the tomato rhizosphere. (A) Nationwide soil pH distribution across China with marked sampling sites used in this study. HLJ: Heilongjiang; HB: Hebei; SX: Shaanxi; SD: Shandong; JS: Jiangsu; AH: Anhui; YN: Yunnan; GX: Guangxi. (B-C) Community composition of bacteria (B) and fungi (C) in bulk soil (left) and tomato rhizosphere (right). (D-E) Non-metric multidimensional scaling (NMDS) plots of bacterial (D) and fungal (E) community structure; circles indicate bulk soil and triangles indicate rhizosphere samples. (F) Tomato growth responses to *Ralstonia* infection across different soil types. (G) Comparison of *Ralstonia* colonization efficiency in bulk soil versus rhizosphere soil under pathogen-invaded and non-invaded conditions across different soil types (left). The accompanying bar plot illustrates the number of *Ralstonia* ASVs detected in the rhizosphere of tomato plants grown in various soil types (right). (H) Plant biomass changes following *Ralstonia* invasion. Data are presented as mean ± SD; statistical significance was assessed using one-way ANOVA with Tukey’s post hoc test (*p* < 0.05). (I) Correlation between *Ralstonia* colonization efficiency in the rhizosphere and changes in plant biomass in response to pathogen invasion. Gray shading indicates 95% confidence interval.

Significant differences in unique bacterial and fungal taxa across soil types (Fig. S2C,D) further underscored the regional specificity of microbiomes. Notably, *Ralstonia* colonization was consistently lower in rhizosphere than in bulk soil, but efficiency varied substantially by soil type (Fig. 1G). This pattern suggests that colonization resistance may reflect functional differences among resident microbial communities, not simply the presence or absence of particular taxa. To validate this, tomato plants were inoculated with *Ralstonia* (Fig. 1F). Infection led to varying degrees of growth inhibition, depending on the soil: GX and HB soils showed severe reductions (∼40% in dry weight), whereas JS soil exhibited no significant impact (Fig. 1F,H; Fig. S1C–I). Despite pathogen exposure, *Ralstonia* abundance remained lower in the rhizosphere, consistent with an active microbiome-mediated barrier. However, colonization varied widely, from 237 ± 101 ASV_2 reads in JS soil to 8,248 ± 3,419 in GX soil—a 35-fold difference (Fig. 1G). This variation was strongly associated with plant performance: *Ralstonia* colonization correlated significantly with dry weight loss in most cases. This association was not observed in HB soil, possibly due to additional environmental factors (Fig. 1I).

To identify the environmental factors shaping microbial assembly and colonization outcomes, we performed Mantel tests using Bray–Curtis dissimilarities. Soil pH and ammonium nitrogen (NH₄⁺-N) emerged as major drivers of bacterial (but not fungal) community structure (Fig. S2E), linking abiotic conditions to microbiome functionality. Lower pH and reduced bacterial α-diversity were associated with higher *Ralstonia* colonization (Fig. S2E–G), consistent with a link between disease suppressiveness, microbial community properties, and environmental context; however, these relationships remain correlative in the present dataset. As a control, *Ralstonia* was cultured in sterile soil extracts. Growth patterns were inversely related to soil pH—highest in JS and lowest in AH soils (Fig. S2H,I). However, in *vitro* growth in sterile soil extracts did not correlate with rhizosphere colonization or disease severity (Fig. S2J–L), indicating that abiotic conditions alone do not explain pathogen suppression.

### Root-associated fatty acids correlate with *Ralstonia* colonization in the tomato rhizosphere

To identify metabolites associated with differential *Ralstonia* colonization, we performed non-targeted metabolomic profiling of root-associated exudates from plants grown in soils representing high (GX), intermediate (HLJ), and low (JS) colonization levels (Fig. 1G). Across all samples, 1,842 metabolites were detected, including 518 lipids and lipid-like molecules, 172 phenylpropanoids and polyketides, and various other compound classes (Fig. S3A). Partial least squares discriminant analysis (PLS-DA) revealed clear metabolic separation among soils, with principal components explaining 34.3% and 23.9% of the variance (Fig. 2A; Fig. S3A). Of the 1,842 metabolites, 992 were differentially abundant among the three rhizosphere types (Fig. 2B; Fig. S3B–D). Compared to HLJ and GX soils, JS soils exhibited a net downregulation of 372 and 144 metabolites, and an upregulation of 107 and 243 metabolites, respectively. Compared to GX soil, HLJ soils showed an upregulation of 514 metabolites and a downregulation of 104 metabolites (Fig. S3B–D; Table S1). This pattern was especially prominent in the lipid fraction. Lipids and lipid-like molecules were least abundant in JS soil—where *Ralstonia* colonization was minimal—and most abundant in GX soil, which supported the highest colonization (Fig. 2C). Across all samples, total lipid abundance was positively correlated with *Ralstonia* colonization efficiency (Fig. 2D; Fig. S3E,F). Among these metabolites, lauric acid (C12:0, LA) showed the strongest positive association with *Ralstonia* colonization across the tested soils (Fig. 2E).

**Fig. 2.**
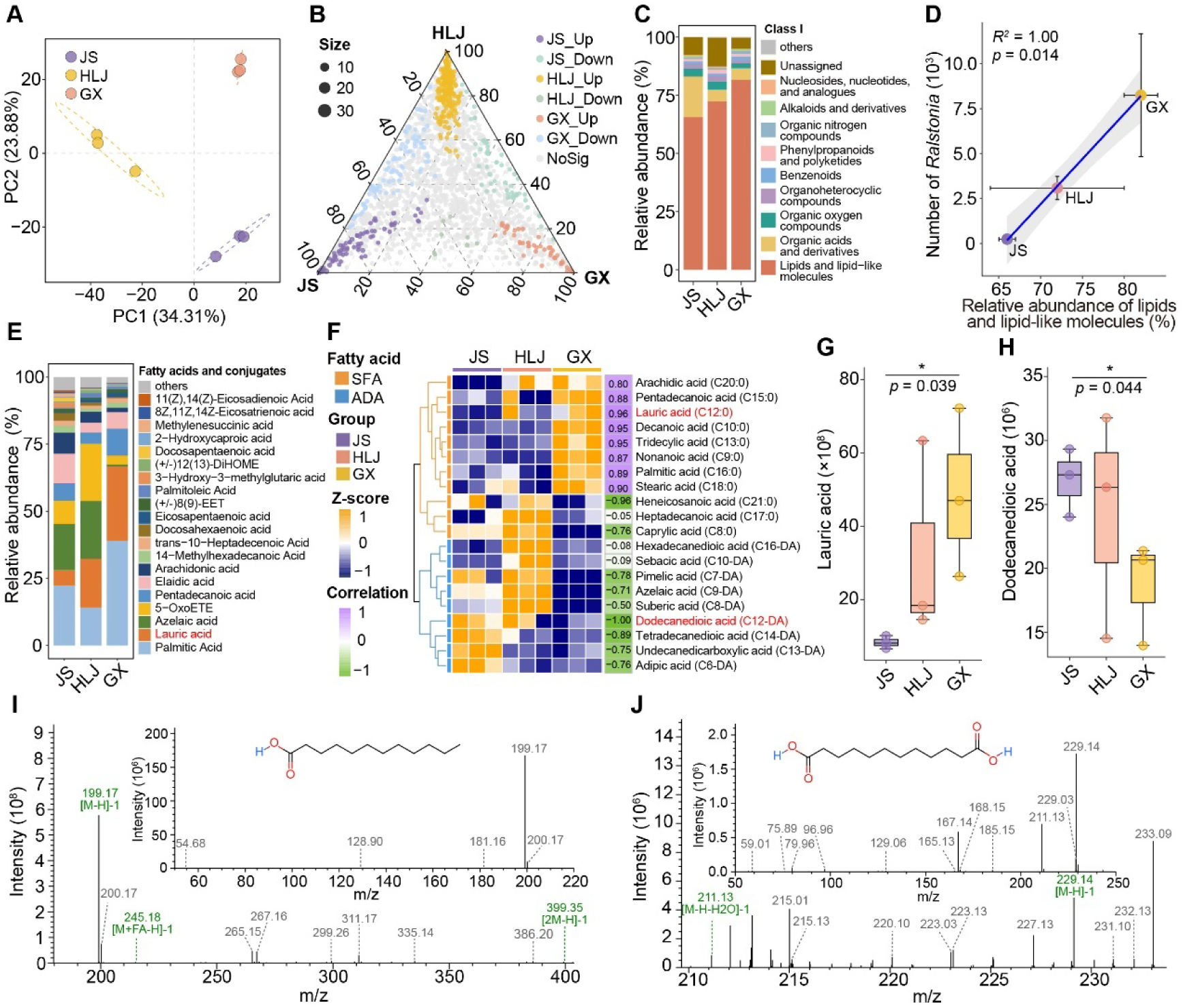
Identification of lauric acid as a key root exudate regulating *Ralstonia* colonization in the tomato rhizosphere. (A) Partial least squares discriminant analysis (PLS-DA) of root exudates from tomatoes grown in different soils. (B) Ternary plots illustrating metabolite profiles identified by non-targeted metabolomics. (C) Relative abundance of major metabolite classes across soil types. (D) Correlation between lipid-related metabolites and *Ralstonia* colonization efficiency. Gray shading represents 95% confidence interval. (E) Relative abundance of fatty acids and derivatives. (F) Abundance and correlation of saturated fatty acids (SFAs) and aliphatic dibasic acids (ADAs) with *Ralstonia* colonization. (G-H) Relative abundance of lauric acid (G) and dodecanedioic acid (H) across soils. (I-J) Identification of lauric acid (I) and dodecanedioic acid (J) by LC-MS/MS. Confirmation using authentic standards is shown in Fig. S3I, J.

Fatty acid profiling revealed that saturated fatty acids (SFAs) such as lauric acid were enriched in GX soils, while aliphatic dibasic acids (ADAs), including dodecanedioic acid (C12-DA), were enriched in JS soils. These two metabolite classes showed opposing correlations with *Ralstonia* colonization (Fig. 2F). Lauric acid and dodecanedioic acid exhibited the strongest contrasting abundance patterns and colonization correlations across soils (Fig. 2F–H), and their identities were validated via LC–MS/MS and authentic standards (Fig. 2I,J; Fig. S3I,J). We quantified the concentrations of individual medium-chain fatty acids (C10–C15) (Fig. S3G). Although several fatty acids were significantly correlated with *Ralstonia* invasion in the rhizosphere, lauric acid was the most abundant species in the rhizosphere of JS soil. Absolute quantification showed that lauric acid levels were approximately 7-fold higher than those of myristic acid, the second most abundant fatty acid within this chain-length class. Lauric acid concentrations increased across compartments more closely associated with the root surface in JS soil, reaching ∼4 μM in bulk soil, ∼33 μM in the rhizosphere, and ∼121 μM in soil tightly adhering to the root surface, while xylem sap contained ∼5 μM LA. Under sterilized conditions, rhizosphere and rhizoplane LA levels increased relative to unsterilized controls, consistent with a microbiome-mediated contribution to LA depletion in these root-associated compartments (Fig. S3H). In GX soil, rhizosphere LA reached ∼120 μM, consistent with weaker buffering of the local LA pool in that background. These measurements define the observed LA pool in each compartment and provide context for microbial exposure.

Together, these findings identify lauric acid as the most experimentally tractable root-associated lipid linked to rhizosphere microbiome composition and pathogen colonization outcomes in this system. We emphasize, however, that additional rhizosphere metabolites also varied across soils and may interact with LA-dependent processes. Although multiple rhizosphere metabolites varied across soils, we focused on lauric acid because it combined four features not shared by most candidates: high abundance in the rhizosphere, strong association with *Ralstonia* colonization across soils, absolute quantifiability across compartments, and clear physiological effects on both pathogen and beneficial taxa.

### Interactions between cultivated rhizobacteria, *Ralstonia*, and lauric acid in the tomato rhizosphere

To characterize the microbial taxa interacting with *Ralstonia* and lauric acid (LA), we employed high-throughput cultivation of rhizosphere bacteria from tomatoes grown in 8 distinct soils (Fig. S1B). This yielded 2,638 bacterial ASVs, dominated by Actinobacteria (37.7%), Proteobacteria (36.4%), Firmicutes (22.5%), and Bacteroidetes (3.2%) (Fig. S4A,B). From these, 1,314 strains were purified, with dominant genera including *Arthrobacter* (29.5%), *Pseudomonas* (16.5%), *Bacillus* (9.6%), and *Microbacterium* (9.6%) (Fig. 3A). To assess antagonism toward *Ralstonia*, all isolates were screened using a double-layer agar assay. Of the 1,314 strains, 248 (18.9%) inhibited *Ralstonia* growth, with antagonistic strains enriched in Proteobacteria (66.1%), followed by Firmicutes (19.0%) and Actinobacteria (14.9%) (Fig. S5A; Fig. 3B,C). Prominent antagonistic genera included *Pseudomonas*, *Bacillus*, and *Arthrobacter*, and the proportion of antagonists across soils correlated with the abundance of *Pseudomonas* and *Bacillus* (Fig. S5A,B). We then evaluated isolate sensitivity to lauric acid, a lipid metabolite enriched in high-*Ralstonia* soils. All 1,314 isolates were exposed to 200 and 500 μM LA (Fig. 3A). Growth was inhibited in 824 isolates (62.7%), with the highest sensitivity in Actinobacteria (75.6%), Firmicutes (73.8%), and Bacteroidetes (72.7%) (Fig. 3D,E). In contrast, only 33.8% of Proteobacteria were sensitive, and 81.8% of LA-tolerant strains belonged to *Pseudomonas* (Fig. S5C). Strikingly, *Bacillus* and *Arthrobacter* were highly susceptible to lauric acid (77.0% and 83.2%, respectively) (Fig. S5C,D). Notably, 18.3% of Firmicutes isolates exhibited both anti-*Ralstonia* activity and LA sensitivity—a vulnerability less common in Proteobacteria (8.8%) and Actinobacteria (3.4%) (Fig. 3F). *Bacillus*, a representative genus within the Firmicutes, was consistently less abundant in rhizosphere samples than in bulk soil, irrespective of pathogen presence (Fig. 3G), supporting the idea that rhizosphere conditions in *vivo* disfavor selected LA-sensitive antagonists. These results identify lauric acid as a microbiome-modulating metabolite associated with pathogen-favoring community shifts, in part by impairing selected beneficial competitors under the tested conditions.

**Fig. 3.**
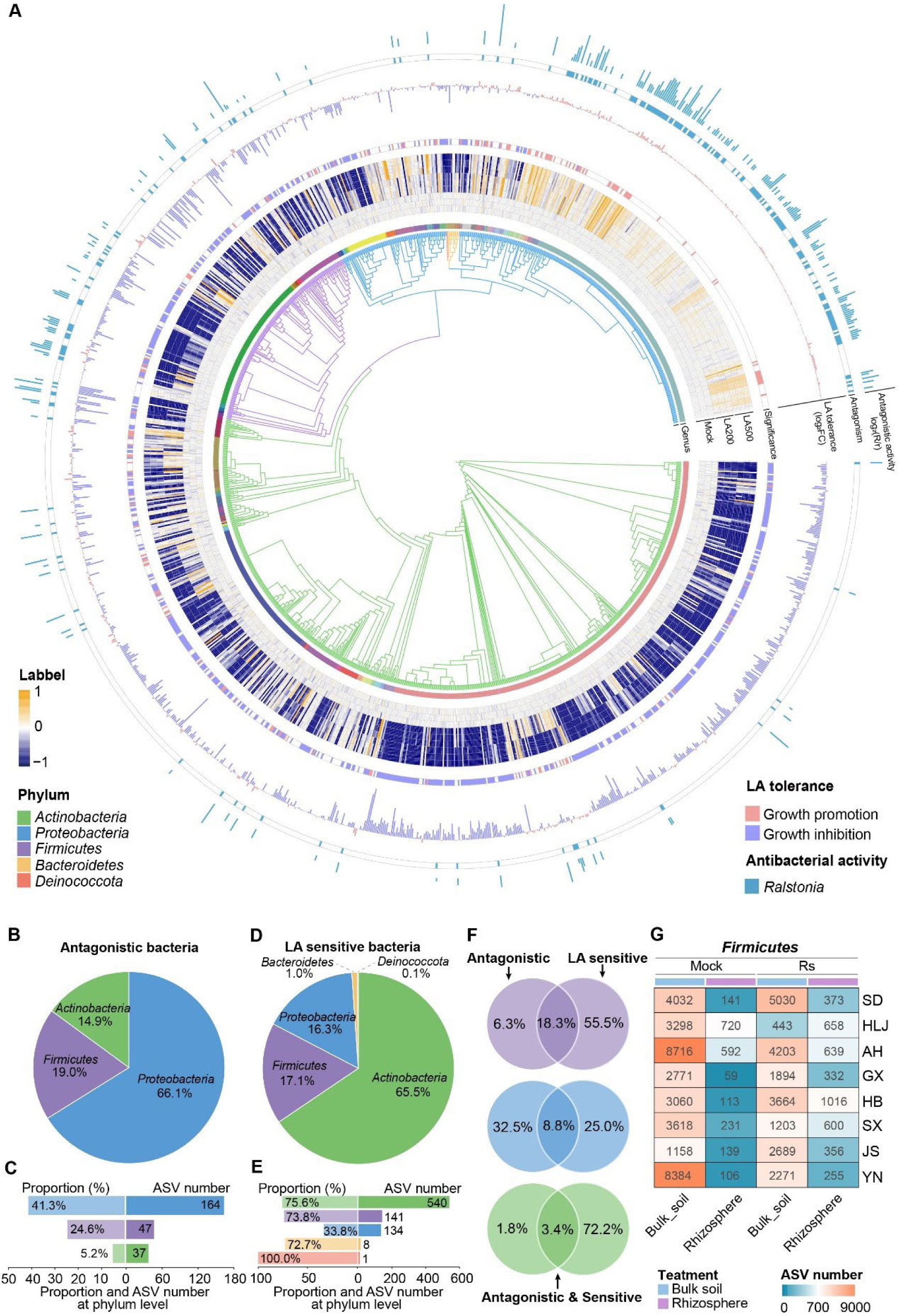
High-throughput cultivation and phenotyping of rhizosphere bacterial communities. (A) Phylogenetic tree of bacterial isolates from tomato rhizosphere. Tree branches are colored by phylum; the inner ring by genus. Heatmap indicates isolate growth at 0, 200, and 500 μM lauric acid. Bar plots show lauric acid tolerance (purple = tolerant; orange = sensitive) and antagonistic effects on *Ralstonia* (Blue). (B) Phylum-level proportions of *Ralstonia*-antagonizing isolates. (C) Number and proportion of antagonistic isolates in Proteobacteria, Firmicutes, and Actinobacteria. (D) Phylum-level distribution of lauric acid-sensitive bacterial isolates, shown as a pie chart. (E) Bar plots showing the proportion and number of ASVs corresponding to lauric acid-sensitive bacterial isolates within the phyla Proteobacteria, Firmicutes, Actinobacteria, Bacteroidetes, and Deinococcota. (F) Proportion of isolates categorized by *Ralstonia* antagonism and lauric acid sensitivity across phyla. (G) Comparison of Firmicutes colonization in bulk and rhizosphere soils, with or without *Ralstonia* invasion.

### Identification of a lauric acid-degrading bacterial community supporting fatty acid homeostasis in the tomato rhizosphere

We therefore asked whether community-level LA turnover could buffer rhizosphere chemistry and reduce disease severity. To test this, we first examined whether LA levels correlate with disease outcomes. In natural soils inoculated with *Ralstonia*, LA amendments caused dose-dependent growth inhibition, with plant height and dry weight reduced by up to 27.1% and 22.5%, respectively (Fig. 4A,B; Fig. S6C). Disease severity also varied by soil: JS soil, which supports strong microbiome-mediated resistance, was least affected, while GX soil was most sensitive. As a control, we repeated the experiment without *Ralstonia* while keeping all other conditions constant. No visible differences in tomato growth were observed between lauric acid–treated and untreated plants. Consistently, quantitative measurements of plant height, fresh weight, and dry weight also showed no statistically significant differences (Fig. S6A,B).

**Fig. 4.**
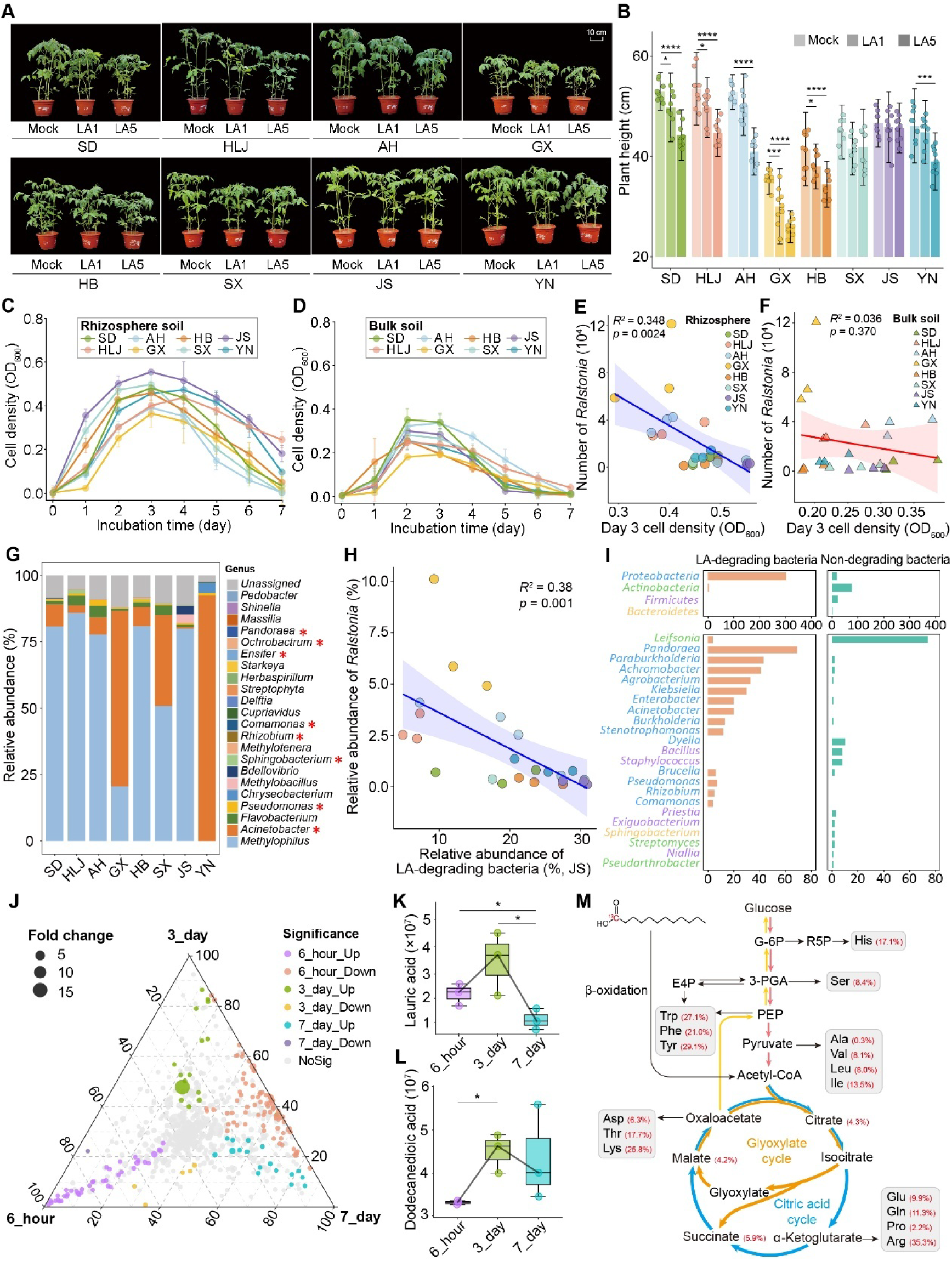
Lauric acid-degrading microbial communities modulate *Ralstonia* colonization resistance. (A-B) Tomato growth (A) and height (B) after *Ralstonia* invasion in soils treated with 0 μM (mock), 50 μM (LA1), or 250 μM (LA5) lauric acid. (C-D) Enrichment of lauric acid-degrading communities in rhizosphere (C) and bulk soil (D) with OD₆₀₀ measured over 7 days. For statistics, see Fig. S6D. (E-F) Correlation between lauric acid degradation activity and *Ralstonia* colonization in rhizosphere (E) and bulk soil (F). (G) Composition of enriched lauric acid-degrading communities with cultivable genera marked with asterisks. (H) Correlation between bacterial species in JS soil-derived LA-degrading communities and *Ralstonia* colonization efficiency. (I) Lauric acid degradation capacity of the isolated bacterial strains. Taxonomic distribution (at the phylum and genus levels) of isolates with (left) and without (right) lauric acid degradation ability. (J) Exometabolomics analysis of culture supernatants at 6 h, 3 d, and 7 d in lauric acid minimal medium. (K-L) Quantification of lauric acid (K) and dodecanedioic acid (L) at each time point. (M) Metabolic flux of ¹³C-labeled lauric acid within lauric-acid-degrading communities. The schematic illustrates the ¹³C labeling ratio of key tricarboxylic acid (TCA) cycle intermediates and amino acids. Specific metabolic pathways are indicated by different colors: TCA cycle (blue), glyoxylate cycle (orange), gluconeogenesis (yellow), and glycolysis (red).

We then enriched LA-degrading communities in minimal medium containing 1 mM LA as the sole carbon source. This concentration was used as a selective enrichment condition to recover microorganisms capable of LA-dependent growth, rather than to mimic the in *situ* concentrations measured in root-associated compartments. Rhizosphere-derived communities exhibited greater LA-degrading capacity than bulk soil counterparts, with JS rhizosphere communities reaching the highest cell densities and GX the lowest (Fig. 4C,D; Fig. S6D). Importantly, enrichment-based LA-degradation capacity of rhizosphere communities showed a strong positive correlation with tomato growth and a strong negative correlation with *Ralstonia* colonization (Fig. 4E; Fig. S6E,F). This association was not observed for bulk-soil communities (Fig. 4F). To further validate the protective effect, we performed in *vitro* infection assays. *Ralstonia* increased disease index by 54.6% in minimal medium, whereas microbial extracts from JS soil reduced severity by 12.1%. LA supplementation (200 μM) increased disease severity by 20.4%, but this effect was counteracted by JS microbial extracts, which lowered disease index by 43.3% (Fig. S6G,H). These findings support a functional association between enrichment-based community-level LA-degrading capacity and disease suppression, which we test further using perturbation, transplantation, and SynCom reconstruction.

Microbial community analysis of LA-degraders across soils revealed 38 genera, mainly from Proteobacteria and Bacteroidetes (Fig. 4G). JS and YN soils contained the most unique LA-degrading taxa, and 15 ASVs were shared across all soils (Fig. S6K). The relative abundance of these LA-degrading taxa negatively correlated with *Ralstonia* colonization (Fig. 4H), consistent with a protective role. From 427 isolated strains, 52.6% were detected via amplicon sequencing (Fig. S6I), and 307 strains (71.9%) could utilize LA as their sole carbon source (Fig. S6J). These degraders were predominantly Proteobacteria, including *Pandoraea*, *Paraburkholderia*, *Achromobacter*, *Agrobacterium*, *Klebsiella*, *Enterobacter*, and *Burkholderia*. In contrast, *Bacillus* and *Leifsonia*, although present in enrichments, could not metabolize LA (Fig. 4I; Fig. S6L). These findings further highlight that not all beneficial microbes can neutralize pathogen-promoting exudates such as LA, underscoring the specialized role of degraders in community-level resistance. Metabolomic profiling of the LA-degrading community from JS soil revealed dynamic changes in exo-metabolites over 6 hours, 3 days, and 7 days in minimal medium (1 mM LA). Of the 696 metabolites detected, 278 changed significantly over time (Fig. 4J; Fig. S7A–E). LA levels initially rose by day 3 and then dropped sharply by day 7 (Fig. 4K), while dodecanedioic acid (C12-DA)—a predicted product of LA ω-oxidation—peaked at day 3 and declined more gradually (Fig. 4L). Together with ^13^C-labeled LA incorporation into central carbon metabolites and amino acids (Fig. 4M), this temporal pattern is consistent with active microbial LA turnover and assimilation rather than simple persistence of the supplied substrate. Consistently, the LA–degrading consortium exhibited high carbon use efficiency, reaching 67.4%–82.4% by day 3 (Fig. S7F), indicating efficient conversion of lauric acid into biomass via central metabolism. We compared the growth dynamics of the highly efficient LA–degrading community (JS_Coms), the less efficient community (GX_Coms), and *Ralstonia* (Rs) alone. JS_Coms exhibited the fastest growth and reached the highest cell density at stationary phase. In contrast, GX_Coms grew more slowly and attained a stationary-phase density approximately 20% lower. Notably, *Ralstonia* displayed the slowest growth (Fig. S7G). KEGG analysis indicated that although *Ralstonia* harbors most components of the fatty-acid degradation pathway, it lacks two key enzymes: acyl-CoA oxidase and long-chain acyl-CoA dehydrogenase (Fig. S7H,I).

### Higher LA-degrading capacity restores resistance to *Ralstonia* in the rhizosphere

To test the functional role of LA-degrading microbiota enriched in the rhizosphere, we disrupted the indigenous community through soil sterilization and assessed the consequences for plant health and pathogen colonization. In JS soil, sterilization markedly increased susceptibility to *Ralstonia*, resulting in significant reductions in plant height (−27.5%), fresh weight (−25.0%), and dry weight (−33.3%) upon infection (Fig. 5A; Fig. S8A). This was accompanied by a 31.4% decrease in the relative abundance of LA-degrading taxa (from 5.4% to 3.7%; Fig. S8C), an ∼80-fold increase in pathogen abundance (Fig. 5C), and an increase of ∼50 μM in rhizospheric lauric acid (Fig. 5D; Fig. S3H). The sterilization comparison is consistent with a microbiome-mediated contribution to LA depletion in root-associated compartments, although the relative contributions of host release, physicochemical partitioning, and microbial turnover remain unresolved. Together, these results indicate that native rhizosphere communities, including LA-degrading taxa, contribute to maintaining lower LA levels and limiting pathogen proliferation in this system. We next tested whether reintroduction of LA-degrading communities could restore disease resistance. Inoculation of GX soil with either an enriched consortium from JS soil (JS_Coms) or a defined synthetic consortium (JS_SynComs) significantly alleviated disease symptoms (Fig. 5B; Fig. S8B) and reduced *Ralstonia* abundance by 55.1% and 75.3%, respectively (Fig. 5E). Successful establishment of the introduced communities was confirmed by increased relative abundance (JS_Coms: 2.4% to 24.9%; JS_SynComs: 2.2% to 43.3%; Fig. S8D), which was associated with a concomitant reduction in rhizospheric LA levels by ∼ 23μM and 32μM respectively (Fig. 5F). These results indicate that LA-degrading communities can alter rhizosphere chemistry and suppress pathogen colonization in this background.

**Fig. 5.**
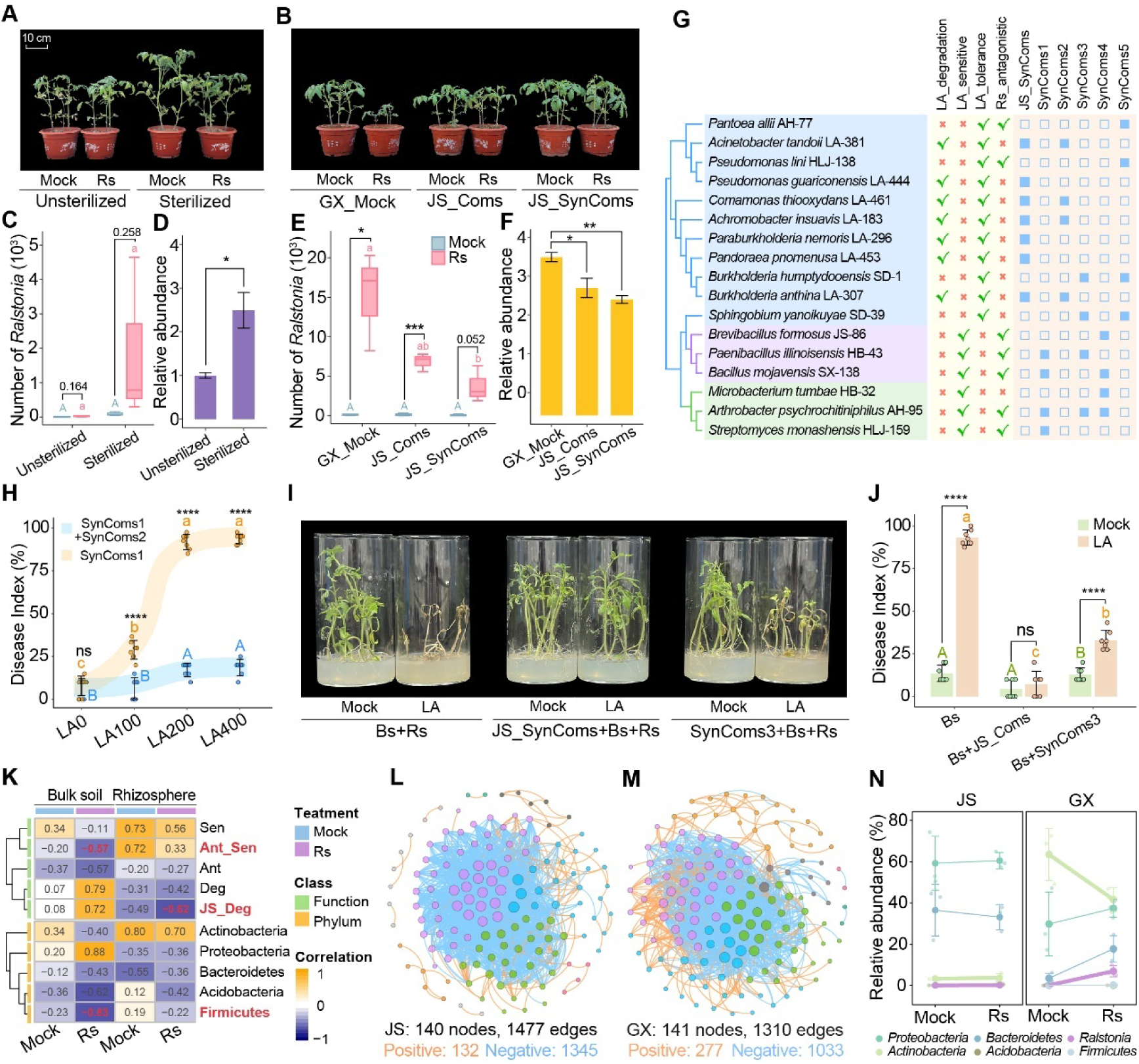
Lauric acid degradation-mediated phenotypic plasticity shapes *Ralstonia* colonization in the tomato rhizosphere. (A) Growth of tomato plants infected with pathogenic *Ralstonia* and grown in unsterilized or sterilized soils from Jiangsu (JS). (B) Growth of infected tomato plants grown in Guangxi (GX) soil supplemented with either a LA-degrading community (JS_Coms) or a synthetic LA-degrading consortium (JS_SynComs). (C and E) Rhizosphere colonization by *Ralstonia* in (C) unsterilized/sterilized JS soil and (E) GX soil under control, JS_Coms, and JS_SynComs treatments. (D and F) Relative abundance of lauric acid in the rhizosphere corresponding to the treatments in (C) and (E), respectively. (G) Schematic representation of the synthetic communities. The phylogenetic tree indicates strain relationships. Functional traits are highlighted in yellow, and strain combinations are shown in orange. JS_SynComs denotes the LA-degrading synthetic community derived from the JS sample. Other consortia include: SynComs1 (antagonistic, LA-sensitive), SynComs2 (LA-degrading), SynComs3 (LA-non-degrading), SynComs4 (LA-sensitive), and SynComs5 (LA-tolerant). (H) Resistance to *Ralstonia* infection and disease incidence conferred by LA-degrading and antagonism-sensitive communities across a gradient of lauric acid concentrations (0, 100, 200, and 400 μM). (I-J) Effects of tripartite interactions among LA-degrading communities, *Bacillus*, and *Ralstonia* on tomato health and disease incidence. (K) Correlations between functional traits and phylum-level microbial community composition with *Ralstonia* colonization in bulk and rhizosphere soils. (L–M) Co-occurrence network analyses of tomato rhizosphere microbiota in JS soil (L) and GX soil (M). (N) Phylum-level community composition and *Ralstonia* dynamics in the tomato rhizosphere under infected and non-infected conditions in JS and GX soils.

To test the proposed mechanism under controlled conditions, we constructed gnotobiotic synthetic communities with defined functional traits (Fig. 5G; Fig. S8E-G). In sterile systems, plants pre-inoculated with an LA-sensitive, *Ralstonia*-antagonistic consortium (SynComs1) exhibited strongly LA-dependent susceptibility: disease incidence increased from 7.9% (control) to 28.9% at 100 μM LA and 91.8% at 200 μM LA. In contrast, co-inoculation with an LA-degrading consortium (SynComs2) suppressed disease incidence to 6.4%–18.6% under the same conditions (Fig. 5H; Fig. S8E). Similar LA-dependent effects were observed using *Bacillus* as a representative LA-sensitive antagonist. To further quantify the contribution of LA degradation, we compared an LA-degrading synthetic consortium (JS_SynComs) with a non-degrading consortium (SynComs3) in a gnotobiotic system containing tomato, *Bacillus*, and *Ralstonia*. In the presence of LA, JS_SynComs reduced disease incidence to 7.1%, whereas SynComs3 reduced it to 33.0%, relative to the *Bacillus* + *Ralstonia* control. In the absence of LA, neither consortium significantly affected disease outcomes (Fig.5 I,J).

Collectively, these results support microbiome-mediated LA turnover as one functional contributor to rhizosphere colonization resistance against *Ralstonia* invasion in this system. We note that the compared consortia differ in community composition beyond LA-degradation capacity; however, the fact that the protective advantage of JS_SynComs emerged specifically under exogenous LA, but not in the no-LA control, argues that LA turnover is a mechanistically important component of the observed phenotype rather than a purely generic community effect.

### Spatial organization of LA-sensitive and LA-degrading microbial taxa across bulk soil and rhizosphere

Across JS, HLJ, and GX soils, Firmicutes, particularly *Bacillus* spp., were consistently enriched in bulk soil, whereas taxa with predicted LA-degrading capacity were associated with rhizosphere Proteobacteria in JS soil and with bulk-soil communities in GX soil (Fig. S8H). These patterns are consistent with spatial structuring of metabolically distinct microbial guilds across soil compartments. Despite its pathogenic potential, *Ralstonia* exhibited limited persistence in the rhizosphere prior to infection, likely reflecting constraints imposed by rhizosphere chemistry and resident microbiota. Following invasion, *Ralstonia* abundance in the rhizosphere was negatively associated with LA-degrading Proteobacteria, suggesting that these taxa restrict pathogen establishment, potentially by reducing local LA availability or altering niche accessibility. In contrast, prior to infection, *Ralstonia* abundance correlated positively with LA-sensitive Actinobacteria, consistent with a permissive niche in the absence of effective metabolic buffering. In bulk soil, *Ralstonia* abundance was negatively correlated with *Bacillus* following infection, supporting a role for Firmicutes in pathogen suppression outside the rhizosphere (Fig. 5K; Fig. S8I,J).

Together, these observations are consistent with spatial segregation of functionally distinct microbial groups. *Bacillus*-dominated communities in bulk soil are associated with antagonism toward *Ralstonia*, whereas rhizosphere-enriched Proteobacteria with LA-degrading capacity are associated with reduced pathogen establishment, plausibly through LA-dependent niche modification. This compartmentalization suggests that invasion outcomes are governed by the balance between LA-sensitive and LA-degrading taxa across soil microenvironments.

Soil-specific differences further modulated these dynamics. GX soil, characterized by elevated rhizosphere LA concentrations and higher abundance of LA-sensitive taxa, exhibited greater susceptibility to LA-driven perturbation and reduced community stability compared to JS soil. Co-occurrence network analysis supported this interpretation, revealing that JS communities displayed fewer positive interactions, more negative associations, and greater network robustness to environmental disturbance (Fig. 5L,M; Fig. S8K,L). Consistently, following *Ralstonia* infection, GX microbiomes underwent more pronounced compositional shifts, including expansion of Proteobacteria—driven in part by *Ralstonia* proliferation—and depletion of LA-sensitive taxa such as Actinobacteria (Fig. 5N; Fig. S8M).

Collectively, these findings support a model in which LA acts as an ecological filter that helps organize microbial communities across soil compartments, with consequences for pathogen invasion and microbiome stability.

### Lauric acid differentially modulates bacterial motility and lysis in *Ralstonia* and *Bacillus*

To investigate how root exudates shape bacterial physiology during the transition from bulk soil to the rhizosphere, we screened 53 representative metabolites—including sugars, amino acids, organic acids, fatty acids, and phenolics—for their effects on bacterial spreading in 3 *Ralstonia* and 4 *Bacillus* strains. Where available, absolute rhizosphere concentrations are provided in Table S2, allowing the screening results to be interpreted relative to measured exposure ranges. Several compounds, including lauric acid, myristic acid, *p*-coumaric acid, and fumaric acid, consistently enhanced *Ralstonia* while suppressing *Bacillus* spreading. In contrast, sugars (e.g., D-galactose, D-fructose, maltose) and select organic acids (e.g., oxalic, lactic, and acetic acid) exhibited the opposite pattern (Fig. S9A). Among these, saturated fatty acids—particularly lauric acid (LA)—elicited the most pronounced genus-specific effects. LA markedly enhanced *Ralstonia* (Fig. 6A,B) while reducing *Bacillus* spreading by 48.7-76.5% (Fig. S9D,E).

**Fig. 6.**
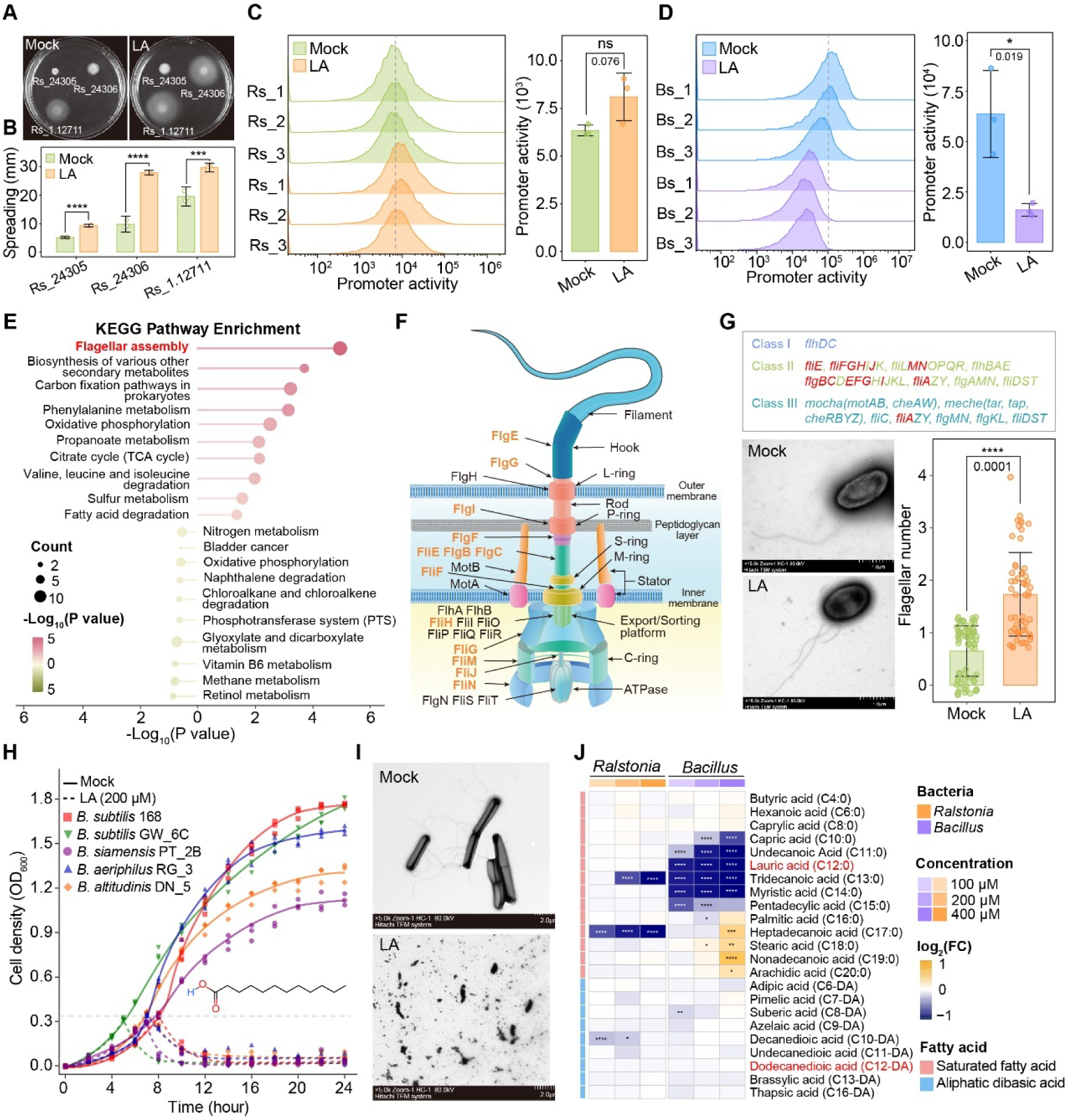
Lauric acid influences the growth and spreading of *Ralstonia* and *Bacillus*. (A-B), Representative images (A) and quantification (B) of colony expansion on soft agar in the presence or absence of lauric acid. (C-D) Flow cytometry quantification of flagellar gene promoter activity: upregulation of *fliC* in *Ralstonia* (C) and repression of *hag* in *Bacillus* (D). (E) KEGG pathway enrichment analysis of differentially expressed genes in *Ralstonia* (red = upregulated; green = downregulated). (F) Schematic of hierarchical regulation of flagellar genes with significantly upregulated genes highlighted in orange. (G) TEM images of *Ralstonia* flagella and quantification of flagellar number with/without lauric acid. (H) Growth of *Bacillus* strains at OD₆₀₀ = 0.3–0.4 with or without 200 μM lauric acid. (I) TEM images showing structural changes in *Bacillus* cells and flagella under lauric acid treatment. (J) Impact of fatty acid chain length, carboxyl group number, and concentration on bacterial growth.

At the transcriptional level, LA increased *Ralstonia P_fliC_*activity by 27.6% (Fig. 6C; Fig. S9B,C), while repressing *Bacillus P_hag_* activity by 74.7% (Fig. 6D; Fig. S9B,C). RNA-seq analysis of LA-treated *Ralstonia* identified 303 upregulated and 226 downregulated genes, with clear separation from controls in PLS-DA space (Fig. S10A). Functional enrichment revealed upregulation of pathways associated with flagellar assembly and the tricarboxylic acid (TCA) cycle. Notably, 14 genes involved in flagellar hook-basal body formation (e.g., *fliA*, *fliG*, *flgE*) and 7 TCA cycle genes (e.g., *sucD*, *sdhA*, *icd*) were significantly induced (Fig. 6E,F; Fig. S10B,C). Transmission electron microscopy (TEM) confirmed a modest increase in flagellar number (∼1.1 additional filaments per cell) in LA-treated *Ralstonia* (Fig. 6G). Functionally, disruption of chemotaxis (*cheA*, *cheY*) or motility (*fliC*, *motA*) genes significantly attenuated *Ralstonia* virulence in tomato (Fig. S10D-F), supporting a link between motility-associated traits and pathogenicity.

In contrast, LA triggered extensive transcriptional reprogramming in *Bacillus* before overt lysis, with 807 genes upregulated and 929 downregulated, whereas the structurally related dodecanedioic acid (C12-DA) elicited comparatively minor changes (Fig. S10G,H). KEGG analysis indicated broad repression of motility-related pathways, including ∼30 flagellar assembly genes and ∼19 chemotaxis genes, alongside induction of glycolysis (∼25 genes). Both LA and C12-DA upregulated non-ribosomal peptide biosynthesis pathways associated with siderophore production (Fig. S10I,J). Consistent with these transcriptional changes, LA induced rapid cell lysis in *Bacillus*, as evidenced by sharp declines in cell density during exponential growth (Fig. 6H) and severe membrane disruption observed by TEM (Fig. 6I; Fig. S10K,L). We also evaluated the growth responses of both *Bacillus* and *Ralstonia* across a concentration gradient of 0–200 μM lauric acid (Fig. S9H-J). Under 40 μM lauric acid treatment—a concentration comparable to that found in the rhizosphere of tomato grown in JS soil—the growth of *Bacillus* was significantly inhibited, with biomass reduced by 9.0% relative to the control. When the concentration was increased to 150 μM, approaching the levels detected near the root surface, the inhibitory effect was substantially enhanced, resulting in a 71.9% reduction in growth. At 200 μM, *Bacillus* was completely unable to grow (Fig. S9F,H,I). In contrast, lauric acid across the entire concentration gradient had no significant effect on the growth of *Ralstonia*, indicating a degree of species-specific susceptibility (Fig. S9F,J).

To assess structure–function relationships, we profiled 14 saturated fatty acids (C4:0–C20:0) and 9 aliphatic dibasic acids (C6-DA–C16-DA). While only a subset inhibited *Ralstonia* growth, six mid-chain fatty acids (C10:0–C15:0) strongly suppressed *Bacillus* viability. In contrast, dibasic acids had minimal effects on *Bacillus* growth (Fig. 6J). Motility assays revealed that *Ralstonia* spreading was selectively enhanced by LA and several dibasic acids, whereas *Bacillus* motility was broadly inhibited by saturated fatty acids, with maximal suppression observed for C11:0–C13:0. Extending this analysis to *Escherichia coli* and *Pseudomonas aeruginosa* revealed additional genus-specific responses, underscoring the importance of fatty acid structure in shaping bacterial behavior (Fig. S9G).

Collectively, these findings identify LA as a rhizosphere-associated metabolite with selective and opposing effects across microbial functional groups. In our assays, LA enhanced *Ralstonia* motility-and virulence-associated gene expression while inhibiting selected LA-sensitive beneficial taxa, particularly Gram-positive antagonists such as *Bacillus*. These results support a model in which disease outcomes depend on the balance between LA-sensitive protective taxa and LA-tolerant or LA-degrading community members. The concentrations used in *vitro* overlapped with those measured across plant-associated compartments, particularly the rhizosphere and root-proximal microhabitats, indicating that the experimental range captures physiologically relevant exposure levels for selected niches. Because the genetic basis of LA biosynthesis and secretion in tomato roots remains unresolved, direct host-side perturbation of LA production was not feasible in this study; accordingly, rhizosphere LA abundance should not be interpreted as a direct proxy for host secretion alone.

## Discussion

Understanding the mechanisms underlying pathogen invasion and microbiome-mediated colonization resistance is critical for the development of sustainable disease management strategies in both plants and animals^73–75^. Our results support a model in which microbiome-mediated turnover of a root-associated fatty acid contributes to rhizosphere colonization resistance against *Ralstonia*^2,68,69^. In this tomato system, protective microbial communities appear to influence pathogen invasion not only through direct antagonism or niche competition, but also by modifying local metabolite availability. This buffering capacity varied across soils and was associated with the balance between LA-sensitive antagonists and LA-tolerant or LA-degrading taxa. We therefore interpret rhizosphere LA abundance as a composite ecological state shaped by plant-derived inputs and microbiome-mediated turnover, rather than as a direct proxy for secretion alone. In this framework, microbiome-mediated fatty-acid turnover emerges as one ecologically grounded mechanism linking local root-associated lipid availability to pathogen invasion outcomes in the tomato rhizosphere (Fig. 7).

**Fig. 7.**
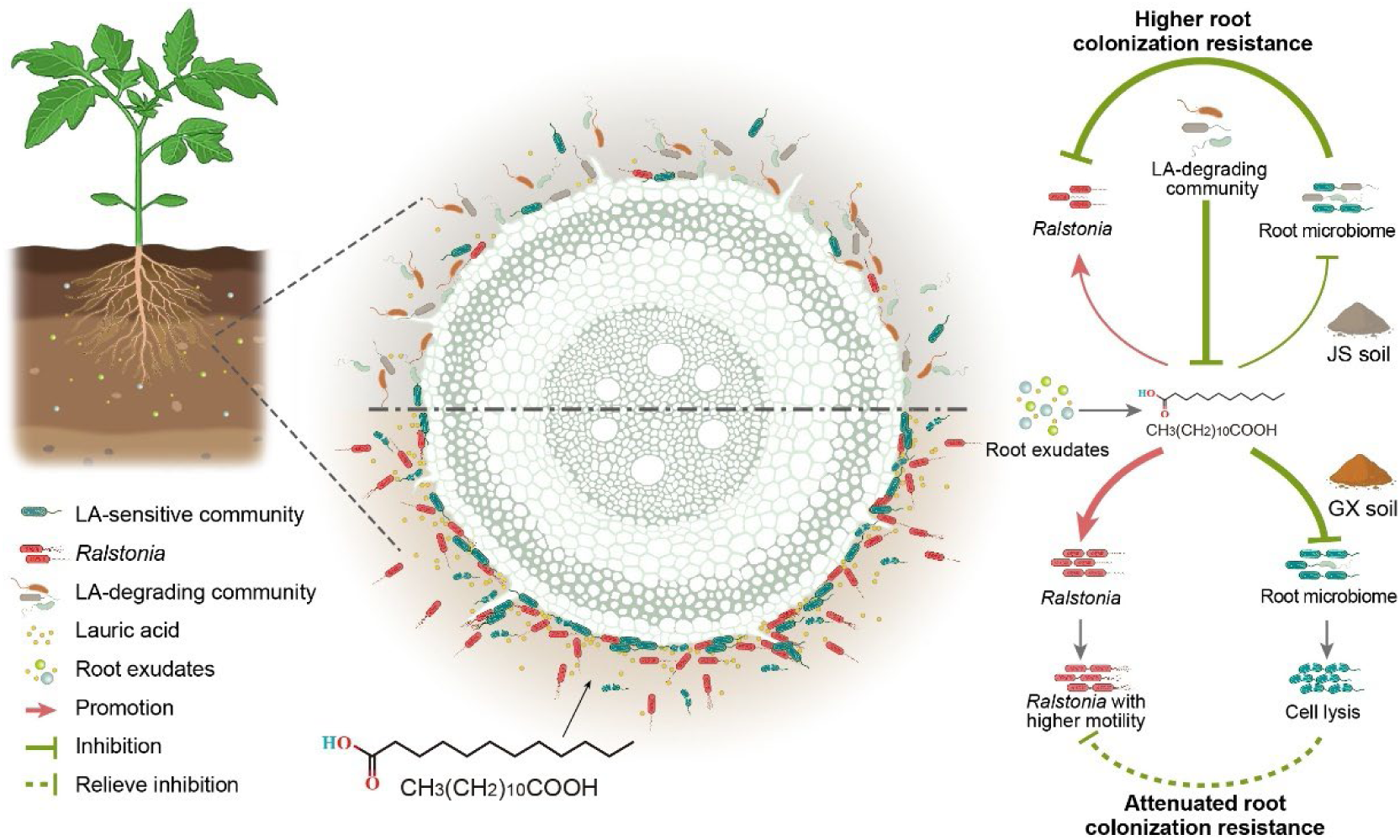
Model for microbiome-mediated lauric acid turnover and colonization resistance in the tomato rhizosphere. Tomato roots are proposed to contribute to the local LA pool at the root–soil interface, where the observed LA distribution is shaped by plant-associated input, physicochemical partitioning, and microbial turnover. During the transition from bulk soil to the rhizosphere, elevated LA enhances *Ralstonia* motility and virulence-associated behaviors while suppressing selected LA-sensitive antagonistic taxa, particularly Gram-positive bacteria like *Bacillus*. In disease-suppressive soils (JS), rhizosphere communities enriched in LA-degrading taxa lower local LA availability and thereby restrict pathogen colonization. In disease-conducive soils (GX), elevated LA is associated with reduced abundance of LA-sensitive taxa, lower microbiome stability, and greater vulnerability to *Ralstonia* invasion. This model further suggests a compartment-specific division of labor in which bulk-soil antagonists and rhizosphere LA-degrading taxa act in complementary ways to promote pathogen exclusion.

### Root microbiome composition and lipid exudates shape disease outcomes

Tomato plants grown in soils with varying characteristics displayed notable differences in growth, disease severity, and *Ralstonia* colonization (Fig. 1). These differences were associated with variation in rhizosphere microbial composition, regional soil properties, and the local lipid environment. Lauric acid emerged as a key root-associated lipid whose local abundance appears to reflect the combined effects of plant-associated input, physicochemical partitioning, and microbiome-mediated turnover. Elevated LA levels were associated with increased *Ralstonia* colonization, enrichment of LA-sensitive taxa in the rhizosphere, and depletion of selected beneficial taxa in bulk soil, including *Bacillus* spp. Lauric acid promoted *Ralstonia* motility- and virulence-associated behaviors while suppressing selected LA-sensitive beneficial taxa, showing how local LA availability can shift the balance between pathogen invasion and microbiome-mediated protection. Soils with greater microbial diversity and higher lipid-degradation capacity showed lower disease severity, whereas soils with reduced bacterial diversity and lower lipid-degradation capacity showed higher *Ralstonia* colonization. Furthermore, specific soil characteristics—such as pH and NH₄⁺-N levels—shaped rhizosphere microbiome composition and lipid degradation potential, indirectly influencing *Ralstonia* colonization. In this tomato–*Ralstonia* system, variation in pathogen colonization was associated not only with host context but also with differences in rhizosphere microbiome composition and putative lipid-degradation capacity^49,76–78^. Although our data identify LA as the most experimentally tractable and ecologically relevant metabolite in this system, we do not suggest that it alone determines pathogen colonization or disease severity. Multiple rhizosphere metabolites varied across soils, and additional lipids or exudate-derived compounds may interact with LA-dependent processes. We focused on LA because it combined high abundance, robust association with colonization outcomes, validated annotation, and clear physiological effects on both pathogen and protective community members.

### Compartment-specific contributions of LA-sensitive antagonists and LA-degrading taxa

Our findings suggest that colonization resistance to *Ralstonia* in this system reflects the interaction of at least two microbial guilds with distinct spatial and functional roles. LA-sensitive antagonists, including many Gram-positive taxa, may contribute most strongly to pathogen suppression in bulk soil or under lower LA exposure. In contrast, LA-tolerant and LA-degrading taxa enriched in the rhizosphere can reduce local LA abundance and help preserve a more protective microbial state near the root surface. Our data support this compartment-specific division of labor, although its full mechanistic basis remains to be resolved. This differential sensitivity and spatial partitioning may drive shifts in microbial community composition in response to root exudates and contribute to the breakdown of colonization resistance in certain soils. Together, these findings highlight a microbial–metabolite interface in which pathogen success is shaped not only by virulence traits but also by metabolite-dependent remodeling of the ecological niche^2,79,80^.

### Lauric acid-degrading consortia reduce local lauric acid abundance and support colonization resistance

Our data suggest that LA-degrading communities can help buffer rhizosphere chemistry during *Ralstonia* invasion. In JS soil, these communities were associated with lower rhizosphere LA abundance and reduced pathogen burden, and transplantation of an LA-degrading community from JS into GX reduced both LA abundance and pathogen colonization (Fig. 5). Microbial conversion of LA into central metabolic intermediates and biomass is consistent with depletion of a metabolite associated with pathogen colonization. These findings support a working model in which antagonistic or niche-overlapping bacteria contribute directly to pathogen exclusion, whereas degradative consortia contribute indirectly by buffering local LA availability at the root–soil interface. The relative contribution of each guild likely depends on compartment, local LA concentration, and community composition. Similar logic has been proposed for host-associated lipids in other systems, but here we support it experimentally only in the tomato rhizosphere^3,81–83^.

### Fatty acids as context-dependent regulators of bacterial physiology in the rhizosphere

A striking finding of this study is the identification of long-chain saturated fatty acids, particularly lauric acid, as key mediators of microbial competition in the rhizosphere (Fig. 2). Rhizosphere compartments with higher LA abundance were associated with increased *Ralstonia* colonization, a higher abundance of LA–sensitive microbial taxa, and a reduced community capacity for lauric acid degradation, particularly in high-disease soils. Functional analyses indicate that lauric acid enhances flagellar assembly- and TCA-associated programs in *Ralstonia*, consistent with improved colonization potential. In contrast, lauric acid downregulated motility-associated programs in *Bacillus* and suppressed growth through envelope damage and lysis in *vitro* (Fig. 6). This differential sensitivity suggests that local lipid availability can contribute to rhizosphere community filtering, thereby influencing niche partitioning between pathogens and beneficial microbes. These findings suggest that metabolite-mediated ecological filtering can shape pathogen invasion alongside, and potentially in interaction with, host immune processes that were not directly dissected here.

Our study highlights fatty-acid-mediated microbial interactions as one contributor to rhizosphere pathogen dynamics. Comparisons with bile acid and fatty acid metabolism in the human gut are conceptually useful because, in both cases, resident microbiota can transform host-associated lipids in ways that alter community structure and pathogen behavior^23,84,85^. However, our data support this mechanism only in the tomato rhizosphere, and we use the gut comparison as a conceptual analogy rather than as evidence for a conserved defense mechanism across kingdoms. Although we observed strong lytic effects in *vitro*, the measured LA concentration in xylem sap was low (∼5.2 μM) and likely non-lethal to *Bacillus* at the bulk scale. Further work will be required to determine whether localized LA accumulation near root surfaces or within tissues can reach biologically relevant lytic thresholds in *vivo*.

## Conclusion and outlook

Soil physicochemical properties and indigenous microbial communities jointly shape root microbiome assembly, generating distinct functional states across soils. In our system, tomato rhizospheres differed in their capacity to process lauric acid, and this variation was associated with differences in colonization resistance. LA acts as a context-dependent ecological modulator at the root–soil interface: under the tested conditions, it promotes *Ralstonia* motility-associated behaviors while inhibiting selected Gram-positive beneficial taxa. In soils enriched in LA-degrading consortia, reduced local LA abundance was associated with stronger microbiome-mediated protection. Our data therefore support an ecological model in which the rhizosphere LA pool is jointly shaped by plant-associated input and microbiome-mediated turnover, and in which disease outcome depends on how that net metabolite state interacts with community composition. We conclude that microbiome-mediated turnover of lauric acid is one ecologically grounded mechanism linking rhizosphere lipid chemistry to pathogen invasion outcomes in tomato.

## Limitations of the study

This study has several limitations. First, we did not directly manipulate host LA biosynthesis or release, because the relevant genetic and transport mechanisms remain insufficiently resolved in tomato. Second, although the measured LA concentrations support physiological relevance in the rhizosphere and near the root surface, the microscale in *vivo* concentrations experienced by individual bacteria remain unknown. Third, our data directly support the proposed mechanism in tomato; however, its generalization to other plant species or to cross-kingdom immune principles requires further validation. Finally, while our perturbation, transplantation, and SynCom experiments support a causal contribution of microbiome-mediated LA turnover, additional host-genetic and spatially resolved studies will be required to define the relative contributions of plant exudation, microbial degradation, and microscale chemical heterogeneity.

## Materials and methods

### Tomato varieties, bacterial strains, and growth conditions

Tomato plants (*Solanum lycopersicum* L. cv. Zhongshu No. 4) were used in all experiments. The bacterial strains and plasmids employed in this study are listed in Supplementary Table S3. *Ralstonia* spp. were cultured at 28°C in liquid TTC medium (10 g·L⁻¹ tryptone, 5 g·L⁻¹ D-glucose, 1 g·L⁻¹ casein hydrolysate, and 0.5 g·L⁻¹ 2,3,5-triphenyltetrazolium chloride). For solid media, 16 g·L⁻¹ agar was added to TTC medium^86^. *Bacillus* spp., *Pseudomonas aeruginosa*, and *Escherichia coli* were grown in Luria–Bertani (LB) medium (10 g·L⁻¹ tryptone, 5 g·L⁻¹ yeast extract, and 10 g·L⁻¹ NaCl). Where appropriate, 50 μg·mL⁻¹ kanamycin was added to the medium.

### Strain and promoter construction

In-frame deletions of *fliC*, *motA*, *cheA*, and *cheY* in *R. solanacearum* strain 1.12711 were generated via homologous recombination using the suicide vector pK18*mobsacB*, as previously described^87^. Briefly, upstream and downstream flanking regions were PCR-amplified using primers listed in Supplementary Table S4 and cloned into pK18*mobsacB*. The resulting constructs were introduced into *Ralstonia* competent cells by electroporation. Double-crossover recombinants were selected on NB medium (10 g·L⁻¹ glucose, 5 g·L⁻¹ peptone, 1 g·L⁻¹ yeast extract, 3 g·L⁻¹ beef extract; pH 7.0) supplemented with 10% sucrose, and verified by PCR. To construct green fluorescent protein (GFP) transcriptional reporters, the promoter region of *fliC* was amplified using P*_fliC_*-F/R primers, and the *gfp* gene was obtained from pUA66 P*_rpsL_*-GFP. The hag promoter was amplified using P*_hag_*-F/R primers, with the *gfp* gene from pGFP4412. The *fliC* and *hag* promoter fragments were fused with *gfp* and cloned into the broad-host-range vectors pBBR1MCS-2 and pHY300PLK, respectively. Reporter plasmids were introduced into *R. solanacearum* strain 1.12711 and *B. subtilis* strain 168 via electroporation for promoter activity assays^88^.

### Soil collection and pot experiments

Soil samples were collected from eight regions in China: Harbin (Heilongjiang), Quzhou (Hebei), Ankang (Shaanxi), Weifang (Shandong), Huai’an (Jiangsu), Lu’an (Anhui), Yuxi (Yunnan), and Guilin (Guangxi). Fresh soils were analyzed for physicochemical properties, including pH, NO₃⁻-N, NH₄⁺-N, available phosphorus (AP), available potassium (AK), and soil organic carbon (SOC). SOC was measured via external heating with potassium dichromate. NH₄⁺-N and NO₃⁻-N were quantified using a flow injection autoanalyzer. AP and AK were determined via flame spectrophotometry and the Mo–Sb colorimetric method, respectively^89^. Soil pH was measured using a calibrated pH electrode. For pot experiments, collected soils were air-dried and sieved through a 5-mm mesh to remove debris. Tomato seeds were germinated for three days before transplanting into pots containing 2 kg of soil (three replicates per treatment). At the 4–5 leaf stage, seedlings were inoculated with 20 mL of *Ralstonia* suspension (1 × 10⁸ CFU·mL⁻¹). Three weeks post-inoculation, plant height, fresh weight, and dry weight were measured. Bulk and rhizosphere soils were collected and stored at –20°C for microbial community profiling. To assess the effect of lauric acid on disease suppression and plant growth, tomatoes were infected with *Ralstonia* (1 × 10⁸ CFU·mL⁻¹), and lauric acid (0, 50, or 250 μM) was applied 7 days before *Ralstonia* infection. Basal growth of tomato plants may vary depending on the time of year.

### Soil microbial community-mediated resistance to *Ralstonia* infection

Surface-sterilized tomato seeds (5% NaClO, 10 min) were germinated in tissue culture bottles. At sowing, 5 mL of soil suspension (1 g fresh soil in 10 mL sterile water) was added. One week later, plants were inoculated with 1 mL of *Ralstonia* suspension (1 × 10^8^ CFU·mL⁻¹). Each treatment included three biological replicates. When necessary, 200 μM lauric acid was added to the medium before solidification. Disease severity was assessed two weeks post-inoculation using a 0–4 rating scale: 0, no symptoms; 1, 1–25% wilted leaves; 2, 26–50%; 3, 51–75%; 4, 76–100%. The disease index (DI) was calculated using the formula: DI = Σ(r × n) / (N × 4) × 100, where r is the disease severity rating, n is the number of plants with that rating, and N is the total number of plants^90^.

### DNA extraction and sequencing

Soil genomic DNA was extracted using a CTAB-based protocol. DNA integrity was confirmed by 1% agarose gel electrophoresis, and concentration was measured using a NanoDrop One spectrophotometer (Thermo Fisher Scientific, Waltham, MA, USA). DNA was diluted to 1 ng·μL⁻¹ in sterile water prior to downstream applications. Bacterial and fungal community compositions were assessed by high-throughput amplicon sequencing on the Illumina MiSeq platform. The V5–V7 hypervariable region of the bacterial 16S rRNA gene was amplified using primers 799F and 1193R^91^, while the ITS1 region of fungal rDNA was amplified using ITS5-1737F and ITS2-2043R primers^92^. PCR reactions (15 μL volume) were performed using Phusion® High-Fidelity PCR Master Mix (New England Biolabs, USA), and amplicons were purified using the Qiagen Gel Extraction Kit (Qiagen, Germany). Sequencing libraries were prepared with the TruSeq® DNA PCR-Free Sample Preparation Kit (Illumina, USA) according to the manufacturer’s instructions. Library quality was assessed using a Qubit 2.0 fluorometer and the Agilent Bioanalyzer 2100. Libraries were sequenced on an Illumina NovaSeq platform, generating 250 bp paired-end reads.

### Non-targeted metabolomic analysis

For non-targeted metabolomic profiling, rhizosphere soil was collected from tomato plants grown in JS, HLJ, and GX soils. Soil tightly adhering to roots was gently brushed off with sterile tools and snap-frozen in liquid nitrogen. To characterize exometabolites produced by a lauric acid-degrading community, cells were inoculated into minimal medium containing: (NH₄)₂SO₄ (1.0 g·L⁻¹), MgSO₄ (0.2 g·L⁻¹), K₂HPO₄ (0.5 g·L⁻¹), NaNO₃ (1.0 g·L⁻¹), CaCl₂ (0.05 g·L⁻¹), FeSO₄ (0.02 g·L⁻¹), and 1 mM lauric acid as the sole carbon source. Cultures were incubated at 28 °C with shaking (180 rpm), and samples were harvested after 6 h, 3 d, and 7 d. Cell suspensions were centrifuged at 12,000 rpm, filtered through 0.2 μm membranes, and snap-frozen in liquid nitrogen. For LC-MS/MS analysis, samples (1 mg·μL⁻¹) were extracted with prechilled 80% methanol, incubated on ice for 5 min, and centrifuged at 15,000 g (4°C, 20 min). Supernatants were diluted with LC-MS–grade water to 53% methanol, centrifuged again, and subjected to analysis^93^. Metabolomic profiling was performed using a Vanquish UHPLC system coupled to an Orbitrap Q Exactive™ HF or HF-X mass spectrometer (Thermo Fisher Scientific) at Novogene Co., Ltd. (Beijing, China). Samples were separated on a Hypersil Gold column (100 × 2.1 mm, 1.9 μm) with a 12 min linear gradient at 0.2 mL·min⁻¹. The mobile phases consisted of eluent A (0.1% formic acid in water) and eluent B (methanol). The gradient program was: 2% B (0–1.5 min), 2–85% B (1.5–3 min), 85–100% B (3–10 min), 100–2% B (10–10.1 min), and 2% B (10.1–12 min). MS parameters included a spray voltage of 3.5 kV, capillary temperature of 320°C, sheath gas flow rate of 35 psi, auxiliary gas flow of 10 L·min⁻¹, S-lens RF level of 60, and auxiliary gas heater temperature of 350°C. Raw data were processed using Compound Discoverer 3.3 (Thermo Fisher) for peak detection, alignment, and quantification. Molecular formulas were predicted based on adducts, precursor ions, and fragmentation patterns. Metabolites were identified by matching against mzCloud, mzVault, and MassList databases. Statistical analyses were performed using R (v3.4.3), Python (v2.7.6), and CentOS (v6.6).

### Transcriptional profiling of bacterial responses to lauric acid

*R. solanacearum* strain 1.12711 was cultured in tryptone broth (5 g·L⁻¹) supplemented with 200 μM lauric acid at 28°C, 180 rpm for 12 h to mid-exponential phase. Methanol-treated cultures served as controls. Cells were harvested by centrifugation at 6,000 rpm, 4°C for 10 min, washed three times with sterile water, frozen in liquid nitrogen, and stored at –80 °C. Each treatment was performed in triplicate. For *B. subtilis* strain 168, cells were grown in TB medium at 28°C, 180 rpm for 4 h, followed by addition of 200 μM lauric acid or dodecanedioic acid. After 1 h incubation, cells were harvested using the same procedure as for *Ralstonia*. Total RNA was extracted and rRNA was removed using targeted probes. RNA was fragmented under elevated temperature with divalent cations. First-strand cDNA synthesis was performed using random hexamer primers, followed by second-strand synthesis using dUTP to maintain strand specificity. Directional libraries were prepared through end-repair, A-tailing, adaptor ligation, size selection, USER enzyme digestion, PCR amplification, and purification. Libraries were diluted to 1.5 ng·μL⁻¹ and validated for insert size distribution. Qualified libraries were pooled based on molarity and sequencing depth requirements, and sequenced on an appropriate platform.

### Promoter activity analysis

The effect of lauric acid on flagellar gene promoter activity in *Ralstonia* and *Bacillus* was evaluated as previously described^94,95^. Strains harboring GFP reporter constructs under the control of the *fliC* (for *Ralstonia*) or *hag* (for *Bacillus*) promoters were grown in tryptone broth supplemented with kanamycin at 28°C with shaking at 180 rpm for 4–6 h. Following incubation with 200 μM lauric acid for 1 h, promoter activity was quantified by measuring GFP fluorescence using flow cytometry (CytoFLEX, Beckman Coulter).

### Bacterial growth analysis

To assess bacterial growth, 1 mL of fresh cell suspension (OD₆₀₀ = 1.0) of *R. solanacearum* strain 1.12711 or *B. subtilis* strain 168 was inoculated into 100 mL of tryptone broth and cultured at 28 °C, 180 rpm. Where indicated, 200 μM lauric acid was added to the culture medium. To evaluate the impact of lauric acid on the growth of various *Bacillus strains* (*B. subtilis* 168, *B. subtilis* GW_6C, *B. siamensis* PT_2B, *B. aeriphilus* RG_3, and *B. altitudinis* DN_5), 1 mL of fresh cell suspension (OD₆₀₀ = 1.0) was inoculated into 100 mL of nutrient broth (NB: glucose 10.0 g·L⁻¹, peptone 5.0 g·L⁻¹, yeast extract 1.0 g·L⁻¹, beef extract 3.0 g·L⁻¹, pH 7.0). Lauric acid (200 μM) was added when cultures reached OD₆₀₀ = 0.3–0.4. Bacteria were grown at 28 °C with shaking (180 rpm) for 24 h. Optical density (OD₆₀₀) was recorded every 2 h. All treatments were performed in triplicate.

### Bacterial response to root exudates

Fifty-three plant-derived metabolites, including 12 carbohydrates, 14 amino acids, 15 organic acids, and 12 phenolic acids, were used to assess their influence on bacterial spreading and gene expression. Three *Ralstonia* strains (CGMCC 1.12711, CICC 24305, CICC 24306) and four *Bacillus* strains (*B. aeriphilus* RG-3, *B. altitudinis* DN-5, *B. siamensis* PT-2B, *B. subtilis* GW-6C) were tested. For spreading assays, 2 μL of cell suspension (OD₆₀₀ = 0.2) was inoculated onto soft agar plates supplemented with 200 μM of each metabolite. Plates were incubated at 28°C for 12 h, and the diameter of bacterial spreading was measured^94^. To evaluate growth and flagellar gene expression in response to metabolites, reporter strains carrying the *fliC* promoter (in *R. solanacearum* 1.12711) or *hag* promoter (in *B. subtilis* 168) were used. A 2 μL aliquot of bacterial suspension (OD₆₀₀ = 0.5) was inoculated into 96-well plates containing 200 μL modified tryptone broth with kanamycin and 200 μM of the indicated metabolite. Cultures were incubated at 28°C with shaking (180 rpm) for 24 h. Cell density (OD₆₀₀) and GFP fluorescence were measured using a Synergy H1 microplate reader (BioTek). All assays were conducted in triplicate.

### High-throughput cultivation and phenotyping of tomato rhizosphere microbiome

High-throughput cultivation of tomato rhizosphere bacteria was performed as previously described^96,97^. Approximately 1 g of rhizosphere soil from eight distinct sites was suspended in 10 mL phosphate-buffered saline (PBS), mixed thoroughly, and serially diluted to 10⁻⁵. Dilutions were inoculated into 96-well plates containing 1/10 tryptic soy broth (TSB) and incubated at 28°C for 2 weeks. Bacterial 16S rRNA gene V5–V7 regions were amplified using primers 799F and 1193R, with dual-barcode PCR for amplicon indexing. Amplicon sequence variants (ASVs) obtained from cultivated bacteria were further isolated and purified on 1/2 strength trypticase soy agar (TSA). Isolate identities were confirmed by 16S rRNA gene sequencing. A total of 1,314 isolates from the eight soils were screened for growth responses to lauric acid in tryptone broth containing 0, 200, or 500 μM lauric acid. Cultures were incubated at 28°C, 180 rpm for 24 h. Antibacterial activity of the isolates against *R. solanacearum* strain 1.12711 was evaluated via plate confrontation assays^98^. Briefly, 1% (v/v) of *Ralstonia* cell culture (OD₆₀₀ = 1.0) was mixed with 200 mL of nutrient agar (glucose 10.0 g·L⁻¹, peptone 5.0 g·L⁻¹, yeast extract 1.0 g·L⁻¹, beef extract 3.0 g·L⁻¹, 1.6% agar, pH 7.0) prior to solidification. Then, 5 μL of bacterial cell suspension (OD₆₀₀ = 1.0) was spotted onto the plates. Inhibition zones were measured after 48 h incubation at 28°C. Antibacterial activity was quantified as: Inhibition index (%) = D/d × 100, where D is the inhibition zone diameter and d is the colony diameter. All experiments were performed in triplicate.

### Enrichment and isolation of lauric acid-degrading bacterial communities

To enrich for LA-degrading community, 1 g of soil was inoculated into 100 mL of minimal medium containing 1 mM lauric acid as the sole carbon source. Cultures were incubated at 28°C, 180 rpm for 72 h, followed by serial re-inoculation into fresh medium for 3–7 enrichment cycles (minimum of three replicates per soil). For isolation of individual strains, enriched cultures were spread on TSB agar (pancreatic digest of casein 17.0 g·L⁻¹, papaic digest of soybean 3.0 g·L⁻¹, dextrose 2.5 g·L⁻¹, NaCl 5.0 g·L⁻¹, K₂HPO₄ 2.5 g·L⁻¹) and incubated. Single colonies were purified through three rounds of streaking and identified by 16S rRNA gene sequencing using primers 27F and 1492R. Lauric acid degradation by each isolate was confirmed in 96-well plates using the same minimal medium. Cell density was measured after 72 h incubation at 28°C, 180 rpm using a microplate reader. To assess differences in LA-degrading capacity among microbial communities from different soils, 1 g of soil was inoculated into 100 mL minimal medium supplemented with 1 mM LA as the sole carbon source. Cultures were incubated at 28°C with shaking at 180 rpm for 7 days, and growth was monitored by measuring cell density at 24 h intervals.

### Construction of synthetic microbial communities (SynComs)

Synthetic microbial communities (SynComs) with distinct functional attributes were assembled from bacterial isolates obtained from the rhizosphere and from the LA-degrading community. Candidate strains were selected on the basis of their LA-degrading capacity, LA tolerance, LA sensitivity, or antagonistic activity. Individual strains were grown overnight in tryptic soy broth (TSB) at 28°C. Cultures were then centrifuged at 5,000 rpm for 5 min, and the resulting cell pellets were washed three times with sterile water. Pellets were resuspended in sterile water and adjusted to an OD_600_ of 0.5 (∼10^8^ CFU·mL^-1^). For SynCom assembly, strains were combined at equal proportions. In total, six SynComs were generated in this study: a simplified SynCom derived from the LA-degrading community (JS_SynComs); an antagonistic and LA-sensitive SynCom (SynComs1); an LA-degrading SynCom (SynComs2); an LA-non-degrading SynCom (SynComs3); an LA-sensitive SynCom (SynComs4); and an LA-insensitive SynCom (SynComs5) (Fig. 5G).

### Microbial growth assays with lauric acid

To assess the effects of lauric acid (LA) on bacterial growth, growth curves of *Ralstonia* and *Bacillus* were determined across a range of LA concentrations. Strains were grown overnight in nutrient broth (NB) at 28°C with shaking at 180 rpm. Cultures were then adjusted to an OD_600_ of 0.5, and 500 μL of each bacterial suspension was inoculated into 50 mL NB medium containing LA at the indicated concentrations (0, 20, 40, 60, 80, 100, 150, or 200 μM). Each treatment was performed with three biological replicates. Cultures were incubated at 28°C with shaking at 180 rpm, and bacterial growth was monitored by measuring OD_600_ every 2 h.

To compare the capacity and efficiency of lauric acid utilization, the high-efficiency LA-degrading community (JS_Coms), the low-efficiency LA-degrading community (GX_Coms), and *Ralstonia* were cultured in minimal medium containing 1 mM LA as the sole carbon source. Cultures were incubated at 28°C with shaking at 180 rpm for 30 h, and samples were collected every 2 h to monitor cell density (OD_600_). Each treatment included three biological replicates. Growth curves were generated from OD_600_ measurements to compare growth dynamics and LA utilization efficiency among the two LA-degrading communities and *Ralstonia*.

### Xylem fluid collection

Xylem fluid was collected from 6-week-old tomato plants (*Solanum lycopersicum* L. cv. Zhongshu No. 4) grown in JS soils under controlled greenhouse conditions. Plants were severed at the stem base using sterile scissors, and the resulting xylem exudate was collected with a sterile syringe and immediately chilled on ice. The collected fluid was centrifuged at 12,000 rpm for 10 min, filter sterilized through a 0.22 μm membrane, and stored at −80°C until use.

### 13C-labeled fatty acid metabolic flux analysis

Microbial samples were extracted with 500 μL of ice-cold extraction solvent (acetonitrile:methanol:water, 4:4:2, v/v/v) and homogenized using a Tissuelyser (JX-24, Jingxin, Shanghai, China) with zirconia beads at 40 Hz for 1 min. Homogenates were centrifuged at 16,000 × g for 15 min at 4°C, and the supernatants were collected and dried under a gentle stream of nitrogen. Dried extracts were reconstituted in 50 μL acetonitrile containing internal standards for UHPLC-HRMS/MS analysis. Extraction was then repeated by adding 1 mL methanol and 1 mL acetonitrile to the remaining material, followed by vortexing, centrifugation, drying, and reconstitution as described above.

UHPLC-HRMS/MS analysis was performed using an Ultimate 3000 UHPLC system (Thermo Fisher Scientific) coupled to a Q Exactive Hybrid Quadrupole-Orbitrap mass spectrometer (Thermo Fisher Scientific). A 2 μL aliquot of each reconstituted extract was injected onto an Acquity UPLC BEH Amide column (100 mm × 2.1 mm, 1.7 μm; Waters) at a flow rate of 0.30 mL·min^-1^. The mobile phases consisted of solvent A (water) and solvent B (90% acetonitrile), each supplemented with 15 mM ammonium acetate (pH 9.0). Metabolites were separated using the following linear gradient: 90% B from 0 to 1 min, decreasing to 75% B at 9 min and 50% B at 10.2 min, held at 50% B until 12 min, then returned to 90% B at 12.1 min and equilibrated until 14 min.

Metabolites were ionized in negative heated electrospray ionization mode with the following source settings: spray voltage, 4.0 kV; capillary temperature, 320°C; probe heater temperature, 320°C; sheath gas, 35 arbitrary units; auxiliary gas, 10 arbitrary units; and S-Lens RF level, 50. Full-scan MS data were acquired at a resolution of 70,000 full width at half maximum (FWHM) at m/z 200 over a mass range of m/z 70–1,050, with an automatic gain control (AGC) target of 3 × 10^6^. Data-dependent acquisition was used to collect MS/MS spectra from the top eight precursor ions in each scan using higher-energy collisional dissociation with normalized collision energies of 15, 30, and 50. MS/MS spectra were acquired at a resolution of 17,500 FWHM with an AGC target of 2 × 10^5^.

### Tomato axenic culture

Seed sterilization was performed as described above. Sterilized tomato seeds were germinated on half-strength Murashige and Skoog (1/2 MS) medium in sterile tissue culture under a 16 h light/8 h dark cycle. Three independent experiments were conducted. In experiment 1, seedlings grown with or without lauric acid (LA; 0, 100, 200, or 400 μM) were inoculated with SynComs1 alone or with a combination of SynComs1 and SynComs2, and were challenged with *Ralstonia* 3 days later. In experiment 2, seedlings grown with or without LA were inoculated with SynComs2, SynComs4, SynComs5, or a combination of SynComs2 and SynComs4, followed by *Ralstonia* challenge 3 days later. In experiment 3, seedlings grown with or without LA were inoculated with *Bacillus* (Bs), JS_SynComs + Bs, or SynComs3 + Bs, and were challenged with *Ralstonia* 3 days after inoculation. Disease incidence was assessed 2 weeks after pathogen inoculation. Each bottle contained 10 seedlings, and each treatment included seven biological replicates.

### Lauric acid-to-biomass conversion efficiency of the LA-degrading community

The conversion efficiency of lauric acid to bacterial biomass was evaluated by measuring the cell dry weight of the LA-degrading community grown with LA as the sole carbon source. Briefly, the LA-degrading community was inoculated at 1% (v/v) into minimal medium (MM) supplemented with 1 mM LA as the sole carbon source. Each treatment was performed with 3 biological replicates. Cultures were incubated at 28°C with shaking at 180 rpm, and samples were collected at 0 h, 6 h, 1 d, 3 d, and 7 d. At each time point, 50 mL of culture was harvested by centrifugation at 7,000 rpm for 10 min at 4°C. Cell pellets were washed three times with sterile water, recollected by centrifugation at 7,000 rpm for 10 min at 4°C, and the supernatants were discarded. The resulting pellets were dried at 80°C to constant weight, and dry biomass was determined using an analytical balance with a precision of 0.0001 g. LA conversion efficiency (*Y_C_*) was calculated as follows:

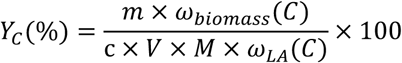

In this equation, *m* denotes the dry cell biomass of the LA-degrading community (mg); *ω_biomass_*(*C*) denotes the carbon content of bacterial biomass, assumed to range from 45% to 55%; *c* denotes the initial concentration of LA (mM); *V* denotes the culture volume (mL); *M* denotes the molar mass of LA (200.32 g mol^-1^); and *ω_LA_*(*C*) denotes the carbon content of LA (71.95%).

### Quantitative metabolomics

Targeted quantitative metabolomics was performed to measure major classes of metabolites commonly found in the rhizosphere, including carbohydrates, amino acids, organic acids, phenolic acids, and fatty acids. For **carbohydrate** analysis, soil samples were extracted with 80% methanol at a 1:10 (w/v) ratio. After thorough homogenization, extracts were incubated at −40°C for 30 min and centrifuged at 15,000 × g for 10 min at 4°C. The supernatant was supplemented with an appropriate volume of a 26-carbohydrate isotope-labeled internal standard mixture (100 μg·mL^-1^; including glucose-^13^C_6_, lyxose-^13^C_6_, ribose-^13^C_6_, and others) and methoxylamine hydrochloride in pyridine (20 mg·mL^-1^). After mixing, samples were incubated at 37°C for 2 h. BSTFA containing 1% TMCS was then added, and samples were incubated at 70°C for 1 h before GC-MS analysis. For **phenolic acid** analysis, soil samples were extracted with 80% methanol containing 1% vitamin C at a 1:2 (w/v) ratio. After grinding and sonication, extracts were centrifuged at 14,000 × g for 10 min at 4°C. The supernatant was passed through a 0.2 μm membrane filter and subjected to LC-MS/MS analysis. For **organic acid** analysis, soil samples were extracted with 80% methanol at a 1:2 (w/v) ratio. Following grinding and sonication, extracts were centrifuged at 14,000 × g for 20 min at 4°C. The supernatant was collected and evaporated to dryness under a gentle stream of nitrogen. Isotope-labeled internal standards (1 μg·mL^-1^ succinic acid-D_4_, 5 μg·mL^-1^ cholic acid-D_4_, and 20 μg·mL^-1^ salicylic acid-D_4_), 50% methanol, and methanol:acetonitrile (1:1, v/v) were then added, and samples were vortexed thoroughly. Derivatization was initiated by adding 3-nitrophenylhydrazine hydrochloride (3-NPH·HCl, 200 mM) and EDC·HCl (120 mM, containing 6% pyridine), followed by mixing and brief centrifugation. Samples were incubated at 40°C for 1 h, vortexed for 30 s, and centrifuged again at 14,000 × g for 20 min at 4°C. The final supernatant was collected for LC-MS/MS analysis. For **amino acid** analysis, soil samples were extracted with acetonitrile:methanol:water (4:4:2, v/v/v) at a 1:2 (w/v) ratio. After sonication, extracts were centrifuged at 16,000 × g for 15 min at 4°C. The supernatant was collected and evaporated to dryness under nitrogen. Sodium carbonate (100 mM) and 2% benzoyl chloride were then added, and derivatization was carried out at room temperature for 10 min. A 21-amino acid isotope-labeled internal standard mixture (1 μg·mL^-1^; including histidine-^13^C_6_, arginine-^13^C_6_, asparagine-^13^C_6_, and others) was subsequently added, and samples were analyzed by LC-MS/MS. For **fatty acid** analysis, soil samples were extracted with acetonitrile:methanol:water (4:4:2, v/v/v) containing C18:0-D_35_ as an internal standard at a 1:5 (w/v) ratio, whereas xylem fluid samples were extracted with the same solvent at a 1:10 ratio. After sonication, extracts were centrifuged at 16,000 × g for 15 min at 4°C. The supernatant was collected and dried under a gentle stream of nitrogen. Residues were reconstituted in 50% methanol and subjected to LC-MS/MS analysis.

### Transmission electron microscopy (TEM) analysis

Flagellar structures in *Ralstonia* were visualized using transmission electron microscopy. Hydrophilized carbon-coated copper grids (400 mesh) were placed at the edge of chemotaxis rings for 2 min to collect cells, followed by brief washes in sterile water and staining with 2% (w/v) uranyl acetate for 5 min. Samples were imaged using a Tecnai Spirit transmission electron microscope operating at 120 kV. For *Bacillus*, *B. subtilis* strain 168 was grown in tryptone broth to OD₆₀₀ = 0.3, then treated with 200 μM lauric acid or methanol (control) for 1 h at 28°C. A 10 μL aliquot of the culture was prepared for TEM as above.

## Statistical analysis

Statistical analyses and visualizations were conducted using GraphPad Prism v8.0 and R (v4.2.1). Differences among treatments were assessed using factorial ANOVA with Tukey’s HSD post hoc tests and Student’s t-tests. Linear regressions were performed using the lm() function from the base R stats package. Correlation plots were generated using the ggpubr package (https://CRAN.R-project.org/package=ggpubr), and all other visualizations were created with ggplot2 in RStudio unless otherwise specified.

## Acknowledgements

This research was supported by the National Key R&D Program of China (2023YFD1900504, 2022YFD1901304), the Chinese Universities Scientific Fund (2022RC033, 2023RC016, 2024RC025), the National Natural Science Foundation of China (32100095) and the Fundamental and the Interdisciplinary Disciplines Breakthrough Plan of the Ministry of Education of China (JYB2025XDXM703, JYB2025XDXM702) to B.N. We thank Prof. Victor Sourjik and Dr. Remy Colin (Max Planck Institute for Terrestrial Microbiology, Marburg, Germany) for their insightful discussions during the revision of this manuscript.

## Author contributions

B.N. designed the research project; S.T., H.L., P.Y., Q.W., T.X., and Y.H. performed the experiments; S.T. wrote the first manuscript; B.N., F. Z. edited the manuscript; and all the authors read and approved the final manuscript.

## Competing interest statement

The authors declare no competing interests.

## Data availability

Bacterial 16S, fungal ITS and RNAseq data for microbial community analysis and transcriptomics in this paper were deposited in the Sequence Read Archive (http://www.ncbi.nlm.nih.gov/sra) under the BioProject ID PRJNA1287328, PRJNA1288679, PRJNA1289397 and PRJNA1438819.

## Notes

### Competing Interest Statement

The authors have declared no competing interest.

## References

1. de la Fuente Canto, C., Simonin, M., King, E., Moulin, L., Bennett, M.J., Castrillo, G., and Laplaze, L. (2020). An extended root phenotype: the rhizosphere, its formation and impacts on plant fitness. Plant J 103, 951–964. 10.1111/tpj.14781.

2. Carrion, V.J., Perez-Jaramillo, J., Cordovez, V., Tracanna, V., de Hollander, M., Ruiz-Buck, D., Mendes, L.W., van Ijcken, W.F.J., Gomez-Exposito, R., Elsayed, S.S., et al. (2019). Pathogen-induced activation of disease-suppressive functions in the endophytic root microbiome. Science 366, 606–612. 10.1126/science.aaw9285.

3. Duran, P., Thiergart, T., Garrido-Oter, R., Agler, M., Kemen, E., Schulze-Lefert, P., and Hacquard, S. (2018). Microbial interkingdom interactions in roots promote *Arabidopsis* survival. Cell 175, 973–983 e914. 10.1016/j.cell.2018.10.020.

4. Zhang, J., Liu, Y.X., Zhang, N., Hu, B., Jin, T., Xu, H., Qin, Y., Yan, P., Zhang, X., Guo, X., et al. (2019). NRT1.1B is associated with root microbiota composition and nitrogen use in field-grown rice. Nat Biotechnol 37, 676–684. 10.1038/s41587-019-0104-4.

5. Bakker, P.A., Pieterse, C.M., de Jonge, R., and Berendsen, R.L. (2018). The soil-borne legacy. Cell 172, 1178–1180. 10.1016/j.cell.2018.02.024.

6. Zhalnina, K., Louie, K.B., Hao, Z., Mansoori, N., da Rocha, U.N., Shi, S., Cho, H., Karaoz, U., Loque, D., Bowen, B.P., et al. (2018). Dynamic root exudate chemistry and microbial substrate preferences drive patterns in rhizosphere microbial community assembly. Nat Microbiol 3, 470–480. 10.1038/s41564-018-0129-3.

7. Loo, E.P., Duran, P., Pang, T.Y., Westhoff, P., Deng, C., Duran, C., Lercher, M., Garrido-Oter, R., and Frommer, W.B. (2024). Sugar transporters spatially organize microbiota colonization along the longitudinal root axis of *Arabidopsis*. Cell Host Microbe 32, 543–556 e6. 10.1016/j.chom.2024.02.014.

8. Cao, Z., Zuo, W., Wang, L., Chen, J., Qu, Z., Jin, F., and Dai, L. (2023). Spatial profiling of microbial communities by sequential FISH with error-robust encoding. Nat Commun 14, 1477. 10.1038/s41467-023-37188-3.

9. Estrela, S., and Brown, S.P. (2013). Metabolic and demographic feedbacks shape the emergent spatial structure and function of microbial communities. PLoS Comput Biol 9, e1003398. 10.1371/journal.pcbi.1003398.

10. Massalha, H., Korenblum, E., Malitsky, S., Shapiro, O.H., and Aharoni, A. (2017). Live imaging of root–bacteria interactions in a microfluidics setup. Proc Natl Acad Sci U S A 114, 4549–4554. 10.1073/pnas.1618584114.

11. Garrido-Sanz, D., and Keel, C. (2025). Seed-borne bacteria drive wheat rhizosphere microbiome assembly via niche partitioning and facilitation. Nat Microbiol 10, 1130–1144. 10.1038/s41564-025-01973-1.

12. Goldford, J.E., Lu, N., Bajic, D., Estrela, S., Tikhonov, M., Sanchez-Gorostiaga, A., Segre, D., Mehta, P., and Sanchez, A. (2018). Emergent simplicity in microbial community assembly. Science 361, 469–474. 10.1126/science.aat1168.

13. He, X., Wang, D., Jiang, Y., Li, M., Delgado-Baquerizo, M., McLaughlin, C., Marcon, C., Guo, L., Baer, M., Moya, Y.A.T., et al. (2024). Heritable microbiome variation is correlated with source environment in locally adapted maize varieties. Nat Plants 10, 598–617. 10.1038/s41477-024-01654-7.

14. Wagner, M.R., Lundberg, D.S., Del Rio, T.G., Tringe, S.G., Dangl, J.L., and Mitchell-Olds, T. (2016). Host genotype and age shape the leaf and root microbiomes of a wild perennial plant. Nat Commun 7, 12151. 10.1038/ncomms12151.

15. Xiong, C., Singh, B.K., He, J.Z., Han, Y.L., Li, P.P., Wan, L.H., Meng, G.Z., Liu, S.Y., Wang, J.T., Wu, C.F., et al. (2021). Plant developmental stage drives the differentiation in ecological role of the maize microbiome. Microbiome 9, 171. 10.1186/s40168-021-01118-6.

16. Fitzpatrick, C.R., Copeland, J., Wang, P.W., Guttman, D.S., Kotanen, P.M., and Johnson, M.T.J. (2018). Assembly and ecological function of the root microbiome across angiosperm plant species. Proc Natl Acad Sci U S A 115, E1157–E1165. 10.1073/pnas.1717617115.

17. Hartman, K., van der Heijden, M.G.A., Wittwer, R.A., Banerjee, S., Walser, J.C., and Schlaeppi, K. (2018). Cropping practices manipulate abundance patterns of root and soil microbiome members paving the way to smart farming. Microbiome 6, 14. 10.1186/s40168-017-0389-9.

18. Sasse, J., Martinoia, E., and Northen, T. (2018). Feed your friends: Do plant exudates shape the root microbiome? Trends Plant Sci 23, 25–41. 10.1016/j.tplants.2017.09.003.

19. Bais, H.P., Weir, T.L., Perry, L.G., Gilroy, S., and Vivanco, J.M. (2006). The role of root exudates in rhizosphere interactions with plants and other organisms. Annu Rev Plant Biol 57, 233–266. 10.1146/annurev.arplant.57.032905.105159.

20. Venturi, V., and Keel, C. (2016). Signaling in the rhizosphere. Trends Plant Sci 21, 187–198. 10.1016/j.tplants.2016.01.005.

21. Badri, D.V., and Vivanco, J.M. (2009). Regulation and function of root exudates. Plant Cell Environ 32, 666–681. 10.1111/j.1365-3040.2008.01926.x.

22. Colin, R., Ni, B., Laganenka, L., and Sourjik, V. (2021). Multiple functions of flagellar motility and chemotaxis in bacterial physiology. FEMS microbiology reviews 45, fuab038. 10.1093/femsre/fuab038.

23. Korenblum, E., Dong, Y., Szymanski, J., Panda, S., Jozwiak, A., Massalha, H., Meir, S., Rogachev, I., and Aharoni, A. (2020). Rhizosphere microbiome mediates systemic root metabolite exudation by root-to-root signaling. Proc Natl Acad Sci U S A 117, 3874–3883. 10.1073/pnas.1912130117.

24. Korenblum, E., Massalha, H., and Aharoni, A. (2022). Plant–microbe interactions in the rhizosphere via a circular metabolic economy. The Plant Cell 34, 3168–3182. 10.1093/plcell/koac163.

25. Macabuhay, A., Arsova, B., Walker, R., Johnson, A., Watt, M., and Roessner, U. (2022). Modulators or facilitators? Roles of lipids in plant root-microbe interactions. Trends Plant Sci 27, 180–190. 10.1016/j.tplants.2021.08.004.

26. da Silva Lima, L., Olivares, F.L., Rodrigues de Oliveira, R., Vega, M.R.G., Aguiar, N.O., and Canellas, L.P. (2014). Root exudate profiling of maize seedlings inoculated with *Herbaspirillum seropedicae* and humic acids. Chem Biol Technol AG 1, 1–18. http://www.chembioagro.com/content/1/1/23.

27. Wippel, K., Tao, K., Niu, Y., Zgadzaj, R., Kiel, N., Guan, R., Dahms, E., Zhang, P., Jensen, D.B., Logemann, E., et al. (2021). Host preference and invasiveness of commensal bacteria in the *Lotus* and *Arabidopsis* root microbiota. Nat Microbiol 6, 1150–1162. 10.1038/s41564-021-00941-9.

28. Erb, M., and Reymond, P. (2019). Molecular interactions between plants and insect herbivores. Annu Rev Plant Biol 70, 527–557. 10.1146/annurev-arplant-050718-095910.

29. Hu, L., Robert, C.A.M., Cadot, S., Zhang, X., Ye, M., Li, B., Manzo, D., Chervet, N., Steinger, T., van der Heijden, M.G.A., et al. (2018). Root exudate metabolites drive plant-soil feedbacks on growth and defense by shaping the rhizosphere microbiota. Nat Commun 9, 2738. 10.1038/s41467-018-05122-7.

30. de Vries, F.T., Griffiths, R.I., Knight, C.G., Nicolitch, O., and Williams, A. (2020). Harnessing rhizosphere microbiomes for drought-resilient crop production. Science 368, 270–274. 10.1126/science.aaz5192.

31. Yu, P., He, X., Baer, M., Beirinckx, S., Tian, T., Moya, Y.A.T., Zhang, X., Deichmann, M., Frey, F.P., Bresgen, V., et al. (2021). Plant flavones enrich rhizosphere Oxalobacteraceae to improve maize performance under nitrogen deprivation. Nat Plants 7, 481–499. 10.1038/s41477-021-00897-y.

32. Stassen, M.J.J., Hsu, S.H., Pieterse, C.M.J., and Stringlis, I.A. (2021). Coumarin communication along the microbiome-root-shoot axis. Trends Plant Sci 26, 169–183. 10.1016/j.tplants.2020.09.008.

33. Thaiss, C.A., Zmora, N., Levy, M., and Elinav, E. (2016). The microbiome and innate immunity. Nature 535, 65–74. 10.1038/nature18847.

34. Teste, F.P., Kardol, P., Turner, B.L., Wardle, D.A., Zemunik, G., Renton, M., and Laliberte, E. (2017). Plant-soil feedback and the maintenance of diversity in Mediterranean-climate shrublands. Science 355, 173–176. 10.1126/science.aai8291.

35. Keppler, A., Roulier, M., Pfeilmeier, S., Petti, G.C., Sintsova, A., Maier, B.A., Bortfeld-Miller, M., Sunagawa, S., Zipfel, C., and Vorholt, J.A. (2025). Plant microbiota feedbacks through dose-responsive expression of general non-self response genes. Nat Plants 11, 74–89. 10.1038/s41477-024-01856-z.

36. Hassani, M.A., Duran, P., and Hacquard, S. (2018). Microbial interactions within the plant holobiont. Microbiome 6, 58. 10.1186/s40168-018-0445-0.

37. Nuccio, E.E., Starr, E., Karaoz, U., Brodie, E.L., Zhou, J., Tringe, S.G., Malmstrom, R.R., Woyke, T., Banfield, J.F., Firestone, M.K., and Pett-Ridge, J. (2020). Niche differentiation is spatially and temporally regulated in the rhizosphere. ISME J 14, 999–1014. 10.1038/s41396-019-0582-x.

38. Gao, M., Teplitski, M., Robinson, J.B., and Bauer, W.D. (2003). Production of substances by *Medicago truncatula* that affect bacterial quorum sensing. Mol Plant Microbe Interact 16, 827–834. 10.1094/mpmi.2003.16.9.827.

39. Zhong, Z., Mu, X., Lang, H., Wang, Y., Jiang, Y., Liu, Y., Zeng, Q., Xia, S., Zhang, B., and Wang, Z. (2024). Gut symbiont-derived anandamide promotes reward learning in honeybees by activating the endocannabinoid pathway. Cell Host Microbe 32, 1944–1958. e7. 10.1016/j.chom.2024.09.013.

40. Takeuchi, T., Kameyama, K., Miyauchi, E., Nakanishi, Y., Kanaya, T., Fujii, T., Kato, T., Sasaki, T., Tachibana, N., and Negishi, H. (2023). Fatty acid overproduction by gut commensal microbiota exacerbates obesity. Cell metabolism 35, 361–375. e9. 10.1016/j.cmet.2022.12.013.

41. Koundouros, N., Nagiec, M.J., Bullen, N., Noch, E.K., Burgos-Barragan, G., Li, Z., He, L., Cho, S., Parang, B., and Leone, D. (2025). Direct sensing of dietary ω-6 linoleic acid through FABP5-mTORC1 signaling. Science 387, eadm9805. 10.1126/science.adm9805.

42. Pascual, G., Domínguez, D., Elosúa-Bayes, M., Beckedorff, F., Laudanna, C., Bigas, C., Douillet, D., Greco, C., Symeonidi, A., and Hernández, I. (2021). Dietary palmitic acid promotes a prometastatic memory via Schwann cells. Nature 599, 485–490. 10.1038/s41586-021-04075-0.

43. Song, X., Zhang, H., Zhang, Y., Goh, B., Bao, B., Mello, S.S., Sun, X., Zheng, W., Gazzaniga, F.S., and Wu, M. (2023). Gut microbial fatty acid isomerization modulates intraepithelial T cells. Nature 619, 837–843. 10.1038/s41586-023-06265-4.

44. Fan, H., Xia, S., Xiang, J., Li, Y., Ross, M.O., Lim, S.A., Yang, F., Tu, J., Xie, L., and Dougherty, U. (2023). *Trans*-vaccenic acid reprograms CD8^+^ T cells and anti-tumour immunity. Nature 623, 1034–1043. 10.1038/s41586-023-06749-3.

45. Oh, S.F., Praveena, T., Song, H., Yoo, J.-S., Jung, D.-J., Erturk-Hasdemir, D., Hwang, Y.S., Lee, C.C., Le Nours, J., and Kim, H. (2021). Host immunomodulatory lipids created by symbionts from dietary amino acids. Nature 600, 302–307. 10.1038/s41586-021-04083-0.

46. Lauson, C.B.N., Tiberti, S., Corsetto, P.A., Conte, F., Tyagi, P., Machwirth, M., Ebert, S., Loffreda, A., Scheller, L., and Sheta, D. (2023). Linoleic acid potentiates CD8^+^ T cell metabolic fitness and antitumor immunity. Cell Metabolism 35, 633–650. e9. 10.1016/j.cmet.2023.02.013.

47. Zhu, M., Frank, M.W., Radka, C.D., Jeanfavre, S., Xu, J., Tse, M.W., Pacheco, J.A., Kim, J.S., Pierce, K., Deik, A., et al. (2024). Vaginal Lactobacillus fatty acid response mechanisms reveal a metabolite-targeted strategy for bacterial vaginosis treatment. Cell 187, 5413–5430.e29. 10.1016/j.cell.2024.07.029.

48. Kachroo, A., and Kachroo, P. (2009). Fatty acid-derived signals in plant defense. Annu Rev Phytopathol 47, 153–176. 10.1146/annurev-phyto-080508-081820.

49. Yuan, J., Zhao, J., Wen, T., Zhao, M., Li, R., Goossens, P., Huang, Q., Bai, Y., Vivanco, J.M., Kowalchuk, G.A., et al. (2018). Root exudates drive the soil-borne legacy of aboveground pathogen infection. Microbiome 6, 156. 10.1186/s40168-018-0537-x.

50. Miao, P., Wang, H., Wang, W., Wang, Z., Ke, H., Cheng, H., Ni, J., Liang, J., Yao, Y.-F., and Wang, J. (2025). A widespread plant defense compound disarms bacterial type III injectisome assembly. Science 387, eads0377. 10.1126/science.ads0377.

51. Jeon, J.E., Kim, J.-G., Fischer, C.R., Mehta, N., Dufour-Schroif, C., Wemmer, K., Mudgett, M.B., and Sattely, E. (2020). A pathogen-responsive gene cluster for highly modified fatty acids in tomato. Cell 180, 176–187. e19. 10.1016/j.cell.2019.11.037

52. Eichmann, R., Richards, L., and Schafer, P. (2021). Hormones as go-betweens in plant microbiome assembly. Plant J 105, 518–541. 10.1111/tpj.15135.

53. Kulkarni, O.S., Mazumder, M., Kini, S., Hill, E.D., Aow, J.S.B., Phua, S.M.L., Elejalde, U., Kjelleberg, S., and Swarup, S. (2024). Volatile methyl jasmonate from roots triggers host-beneficial soil microbiome biofilms. Nat Chem Biol 20, 473–483. 10.1038/s41589-023-01462-8.

54. Kengmo Tchoupa, A., Eijkelkamp, B.A., and Peschel, A. (2022). Bacterial adaptation strategies to host-derived fatty acids. Trends Microbiol 30, 241–253. 10.1016/j.tim.2021.06.002.

55. Levy, M., Kolodziejczyk, A.A., Thaiss, C.A., and Elinav, E. (2017). Dysbiosis and the immune system. Nat Rev Immunol 17, 219–232. 10.1038/nri.2017.7.

56. Takeuchi, T., Kameyama, K., Miyauchi, E., Nakanishi, Y., Kanaya, T., Fujii, T., Kato, T., Sasaki, T., Tachibana, N., Negishi, H., et al. (2023). Fatty acid overproduction by gut commensal microbiota exacerbates obesity. Cell Metab 35, 361–375. e9. 10.1016/j.cmet.2022.12.013.

57. Zhong, Z., Mu, X., Lang, H., Wang, Y., Jiang, Y., Liu, Y., Zeng, Q., Xia, S., Zhang, B., Wang, Z., et al. (2024). Gut symbiont-derived anandamide promotes reward learning in honeybees by activating the endocannabinoid pathway. Cell Host Microbe 32, 1944–1958. e7. 10.1016/j.chom.2024.09.013.

58. Ma, C., Vasu, R., and Zhang, H. (2019). The role of long-chain fatty acids in inflammatory bowel disease. Mediators Inflamm 2019, 8495913. 10.1155/2019/8495913.

59. Tsenkova, M., Brauer, M., Pozdeev, V.I., Kasakin, M., Busi, S.B., Schmoetten, M., Cheung, D., Meyers, M., Rodriguez, F., Gaigneaux, A., et al. (2025). Ketogenic diet suppresses colorectal cancer through the gut microbiome long chain fatty acid stearate. Nat Commun 16, 1792. 10.1038/s41467-025-56678-0.

60. Phair, K., Culliton, D., Kealey, C., and Brady, D. (2024). Long chain unsaturated fatty acids alter growth and reduce biofilm formation of *Cronobacter sakazakii*. J Food Safety 44, e13130. 10.1111/jfs.13130

61. Nakatsuji, T., Kao, M.C., Fang, J.Y., Zouboulis, C.C., Zhang, L., Gallo, R.L., and Huang, C.M. (2009). Antimicrobial property of lauric acid against *Propionibacterium Acnes*: its therapeutic potential for inflammatory acne vulgaris. J Invest Dermatol 129, 2480–2488. 10.1038/jid.2009.93.

62. Genin, S., and Denny, T.P. (2012). Pathogenomics of the *Ralstonia solanacearum* species complex. Annu Rev Phytopathol 50, 67–89. 10.1146/annurev-phyto-081211-173000.

63. Lowe-Power, T.M., Khokhani, D., and Allen, C. (2018). How *Ralstonia solanacearum* exploits and thrives in the flowing plant xylem environment. Trends Microbiol 26, 929–942. 10.1016/j.tim.2018.06.002.

64. Tans-Kersten, J., Huang, H., and Allen, C. (2001). *Ralstonia solanacearum* needs motility for invasive virulence on tomato. J Bacteriol 183, 3597–3605. 10.1128/jb.183.12.3597-3605.2001.

65. Yao, J., and Allen, C. (2006). Chemotaxis is required for virulence and competitive fitness of the bacterial wilt pathogen *Ralstonia solanacearum*. J Bacteriol 188, 3697–3708. 10.1128/jb.188.10.3697-3708.2006.

66. Kloepper, J.W., Ryu, C.M., and Zhang, S. (2004). Induced systemic resistance and promotion of plant growth by *Bacillus* spp. Phytopathology 94, 1259–1266. 10.1094/phyto.2004.94.11.1259.

67. Chen, T., Nomura, K., Wang, X., Sohrabi, R., Xu, J., Yao, L., Paasch, B.C., Ma, L., Kremer, J., Cheng, Y., et al. (2020). A plant genetic network for preventing dysbiosis in the phyllosphere. Nature 580, 653–657. 10.1038/s41586-020-2185-0.

68. Mendes, R., Kruijt, M., De Bruijn, I., Dekkers, E., Van Der Voort, M., Schneider, J.H., Piceno, Y.M., DeSantis, T.Z., Andersen, G.L., and Bakker, P.A. (2011). Deciphering the rhizosphere microbiome for disease-suppressive bacteria. Science 332, 1097–1100. 10.1126/science.1203980

69. Kwak, M.-J., Kong, H.G., Choi, K., Kwon, S.-K., Song, J.Y., Lee, J., Lee, P.A., Choi, S.Y., Seo, M., and Lee, H.J. (2018). Rhizosphere microbiome structure alters to enable wilt resistance in tomato. Nat Biotechnol 36, 1100–1109. 10.1038/nbt.4232

70. Liao, J., Li, Z., Xiong, D., Shen, D., Wang, L., Lin, L., Shao, X., Liao, L., Li, P., Zhang, L.Q., et al. (2023). Quorum quenching by a type IVA secretion system effector. ISME J 17, 1564–1577. 10.1038/s41396-023-01457-2.

71. Grandclement, C., Tannieres, M., Morera, S., Dessaux, Y., and Faure, D. (2016). Quorum quenching: role in nature and applied developments. FEMS Microbiol Rev 40, 86–116. 10.1093/femsre/fuv038.

72. Caballero-Flores, G., Pickard, J.M., and Nunez, G. (2023). Microbiota-mediated colonization resistance: mechanisms and regulation. Nat Rev Microbiol 21, 347–360. 10.1038/s41579-022-00833-7.

73. Singh, B.K., Delgado-Baquerizo, M., Egidi, E., Guirado, E., Leach, J.E., Liu, H., and Trivedi, P. (2023). Climate change impacts on plant pathogens, food security and paths forward. Nat Rev Microbiol 21, 640–656. 10.1038/s41579-023-00900-7.

74. Fitzpatrick, C.R., Salas-Gonzalez, I., Conway, J.M., Finkel, O.M., Gilbert, S., Russ, D., Teixeira, P., and Dangl, J.L. (2020). The plant microbiome: From ecology to reductionism and beyond. Annu Rev Microbiol 74, 81–100. 10.1146/annurev-micro-022620-014327.

75. Huang, Y.X., Tang, S.Y., Liu, R.M., Yu, P.G., Liu, J., Xiao, T., Zhang, Y., Fan, M.S., Zhang, F.S., and Ni, B. (2025). Soil organic carbon mediates plant immunityrhizosphere microbiome interactions and controls colonization resistance to microbial inoculants. Cell Host Microbe 33, 1929–1944. 10.1016/j.chom.2025.10.002.

76. Bakker, P., Berendsen, R.L., Van Pelt, J.A., Vismans, G., Yu, K., Li, E., Van Bentum, S., Poppeliers, S.W.M., Sanchez Gil, J.J., Zhang, H., et al. (2020). The soil-borne identity and microbiome-assisted agriculture: Looking back to the future. Mol Plant 13, 1394–1401. 10.1016/j.molp.2020.09.017.

77. Wei, Z., Gu, Y., Friman, V.P., Kowalchuk, G.A., Xu, Y., Shen, Q., and Jousset, A. (2019). Initial soil microbiome composition and functioning predetermine future plant health. Sci Adv 5, eaaw0759. 10.1126/sciadv.aaw0759.

78. Berendsen, R.L., Pieterse, C.M., and Bakker, P.A. (2012). The rhizosphere microbiome and plant health. Trends Plant Sci 17, 478–486. 10.1016/j.tplants.2012.04.001.

79. Lamichhane, J.R., Barbetti, M.J., Chilvers, M.I., Pandey, A.K., and Steinberg, C. (2024). Exploiting root exudates to manage soil-borne disease complexes in a changing climate. Trends Microbiol 32, 27–37. 10.1016/j.tim.2023.07.011.

80. Jones, J.D.G., Staskawicz, B.J., and Dangl, J.L. (2024). The plant immune system: From discovery to deployment. Cell 187, 2095–2116. 10.1016/j.cell.2024.03.045.

81. Chapelle, E., Mendes, R., Bakker, P.A., and Raaijmakers, J.M. (2016). Fungal invasion of the rhizosphere microbiome. ISME J 10, 265–268. 10.1038/ismej.2015.82.

82. Winston, J.A., and Theriot, C.M. (2020). Diversification of host bile acids by members of the gut microbiota. Gut Microbes 11, 158–171. 10.1080/19490976.2019.1674124.

83. Lee, M.H., Nuccio, S.P., Mohanty, I., Hagey, L.R., Dorrestein, P.C., Chu, H., and Raffatellu, M. (2024). How bile acids and the microbiota interact to shape host immunity. Nat Rev Immunol 24, 798–809. 10.1038/s41577-024-01057-x.

84. Voges, M., Bai, Y., Schulze-Lefert, P., and Sattely, E.S. (2019). Plant-derived coumarins shape the composition of an *Arabidopsis* synthetic root microbiome. Proc Natl Acad Sci U S A 116, 12558–12565. 10.1073/pnas.1820691116.

85. Buffie, C.G., Bucci, V., Stein, R.R., McKenney, P.T., Ling, L., Gobourne, A., No, D., Liu, H., Kinnebrew, M., Viale, A., et al. (2015). Precision microbiome reconstitution restores bile acid mediated resistance to *Clostridium difficile*. Nature 517, 205–208. 10.1038/nature13828.

86. Song, S., Yin, W., Sun, X., Cui, B., Huang, L., Li, P., Yang, L., Zhou, J., and Deng, Y. (2020). Anthranilic acid from *Ralstonia solanacearum* plays dual roles in intraspecies signalling and inter-kingdom communication. The ISME Journal 14, 2248–2260. 10.1038/s41396-020-0682-7.

87. Schäfer, A., Tauch, A., Jäger, W., Kalinowski, J., Thierbach, G., and Pühler, A. (1994). Small mobilizable multi-purpose cloning vectors derived from the *Escherichia coli* plasmids pK18 and pK19: selection of defined deletions in the chromosome of *Corynebacterium glutamicum*. Gene 145, 69–73. 10.1016/0378-1119(94)90324-7.

88. Perrier, A., Barberis, P., and Genin, S. (2018). Introduction of genetic material *in Ralstonia solanacearum* through natural transformation and conjugation. Methods Mol Biol 1734, 201–207. 10.1007/978-1-4939-7604-1_16

89. Geng, J., Yang, X., Huo, X., Chen, J., Lei, S., Li, H., Lang, Y., and Liu, Q. (2020). Determination of the best controlled-release potassium chloride and fulvic acid rates for an optimum cotton yield and soil available potassium. Front Plant Sci 11, 562335. 10.3389/fpls.2020.562335

90. Li, X., Yin, W., Lin, J.D., Zhang, Y., Guo, Q., Wang, G., Chen, X., Cui, B., Wang, M., Chen, M., et al. (2023). Regulation of the physiology and virulence of *Ralstonia solanacearum* by the second messenger 2 ′ ,3 ′-cyclic guanosine monophosphate. Nat Commun *14*, 7654. 10.1038/s41467-023-43461-2.

91. Seybold, H., Demetrowitsch, T.J., Hassani, M.A., Szymczak, S., Reim, E., Haueisen, J., Lübbers, L., Rühlemann, M., Franke, A., Schwarz, K., and Stukenbrock, E.H. (2020). A fungal pathogen induces systemic susceptibility and systemic shifts in wheat metabolome and microbiome composition. Nat Commun 11, 1910. 10.1038/s41467-020-15633-x.

92. Zhang, T., Wang, Z., Lv, X., Li, Y., and Zhuang, L. (2019). High-throughput sequencing reveals the diversity and community structure of rhizosphere fungi of *Ferula Sinkiangensis* at different soil depths. Sci Rep 9, 6558. 10.1038/s41598-019-43110-z.

93. Want, E.J., Masson, P., Michopoulos, F., Wilson, I.D., Theodoridis, G., Plumb, R.S., Shockcor, J., Loftus, N., Holmes, E., and Nicholson, J.K. (2013). Global metabolic profiling of animal and human tissues via UPLC-MS. Nat Protoc 8, 17–32. 10.1038/nprot.2012.135.

94. Ni, B., Ghosh, B., Paldy, F.S., Colin, R., Heimerl, T., and Sourjik, V. (2017). Evolutionary remodeling of bacterial motility checkpoint control. Cell Rep 18, 866–877. 10.1016/j.celrep.2016.12.088.

95. Ni, B., Colin, R., Link, H., Endres, R.G., and Sourjik, V. (2020). Growth-rate dependent resource investment in bacterial motile behavior quantitatively follows potential benefit of chemotaxis. Proc Natl Acad Sci U S A 117, 595–601. 10.1073/pnas.1910849117.

96. Zhang, J., Liu, Y.-X., Guo, X., Qin, Y., Garrido-Oter, R., Schulze-Lefert, P., and Bai, Y. (2021). High-throughput cultivation and identification of bacteria from the plant root microbiota. Nat Protoc 16, 988–1012. 10.1038/s41596-020-00444-7.

97. Huang, Y., Wu, Q., Liu, C., Yu, P., Li, H., Tang, S., Zhang, F., Liu, S., and Ni, B. (2025). Ribosomal RNA operon copy number: A trait-informed framework to close the microbial cultivation gap. iMetaOmics, e70071. 10.1002/imo2.70071.

98. Cao, Y., Pi, H., Chandrangsu, P., Li, Y., Wang, Y., Zhou, H., Xiong, H., Helmann, J.D., and Cai, Y. (2018). Antagonism of two plant-growth promoting *Bacillus velezensis* isolates against *Ralstonia solanacearum* and *Fusarium oxysporum*. Sci Rep 8, 4360. 10.1038/s41598-018-22782-z.

